## Supplementary Material for "Dynamic cybergenetic control of bacterial co-culture composition via optogenetic feedback"

**This PDF file includes:**

Supplementary text section 1-2

Supplementary Figures 1-21

Supplementary table 1

### 1 Formulation of the mathematical modeling framework

#### Unit conversion

We note that the cellular protein density of *E. coli* is approximately independent of the growth rate(4). Because of this, we can estimate a factor that converts protein concentrations into fractions of the proteome and viceversa. We define the proteome fraction of a given protein type $X$ as

$$\Phi_X = \frac{M_X[\mathrm{aa}]}{M_{cell}[\mathrm{aa}]} \tag{1}$$

where $M_X$ is the mass of all protein copies of type $X$ and $M_{cell}$ is the mass of all proteins in the cell, both in units of the average mass of an amino acid. Denoting the mass of a single copy of $X$ by $n_X$ and the number of protein copies by $N_X$, we can relate $\Phi_X$ to the concentration $c_X$ in the following way:

$$\begin{aligned}\Phi_X &= \frac{M_X[\mathrm{aa}]}{M_{cell}[\mathrm{aa}]} = n_X[\mathrm{aa}]\,\frac{N_X[\mathrm{molecules}]}{M_{cell}[\mathrm{aa}]}\\ &= n_X[\mathrm{aa}]\,\frac{V_{cell}[\mathrm{fL}]}{M_{cell}[\mathrm{aa}]}\,\frac{N_X[\mathrm{molecules}]}{V_{cell}[\mathrm{fL}]}\\ &= n_X[\mathrm{aa}]\,\rho_{cell}^{-1}\,c_X[\frac{\mathrm{molecules}}{\mathrm{fL}}]\end{aligned} \tag{2}$$

where we have denoted the cell volume by $V_{cell}$ and introduced the cellular protein density, which is approximately(5) $\rho_{cell} \approx \frac{2\cdot 10^9\,\mathrm{aa}}{1\,\mathrm{fL}} = 2\cdot 10^9\,\frac{\mathrm{aa}}{\mathrm{fL}}$. In the following sections, we use $\frac{n_X}{\rho_{cell}}$ as an inter-conversion factor between protein concentrations and proteome mass fractions.

#### Proteome partition model of bacterial physiology

The framework for host-aware modeling of synthetic genetic circuits we present in this study is based on the proteome-partition model put forward by Scott *et al.*(2). In this section, we provide a brief summary of their model, focusing on the equations that are central to the development of our approach. For further details and derivations of these equations from first principles, we refer the reader to the original publications (2, 3). For examples on how these types of models have been applied to a range of biological problems, see e.g. (6–8).

The model proposed by Scott *et al.* attempts to explain key observations about how the macromolecular composition of *E. coli* varies when the growth rate of the cell changes. The composition of the cell, i.e. the relative amounts of its component proteins, is dictated by its gene expression profile. Therefore, the relation between the composition and the growth rate is ultimately a connection between gene expression and the physiology of the cell, for which growth rate is the most direct readout. The laws governing this relation are of high importance for the prediction of how synthetic genetic circuits behave in the cellular context.

Despite the complexity of a cell, the empirical link between aspects of its macromolecular composition and its growth rate turns out to be remarkably simple. At the core of the model of Scott *et al.* are two linear relations that were determined experimentally, which relate the fraction of the proteome made up by ribosomes and the growth rate (depicted schematically in Figure 4A). The first one applies when the growth rate is changed by modulating the quality of nutrients in the media and it implies a positive correlation between ribosomal content and growth rate:

$$\lambda = \gamma_0\,(\Phi_R - \Phi_{R_0}) \tag{3}$$

where $\lambda$ represents the growth rate, $\Phi_R$, the ribosomal mass fraction and $\Phi_{R_0}$, the Y-offset extrapolated from experiments which corresponds to ribosomes that are not actively engaged in translation. The proportionality constant $\gamma_0$ can be shown to correspond to the translation rate.

The second linear relation is revealed when growth rate is modulated by translation inhibition, e.g. by exposing *E. coli* to increasing concentrations of chloramphenicol in the presence of saturating nutrient availability. In that case, the cell upregulates its ribosome content at lower growth rates, in response to the increasing inhibition of translation brought about by the antibiotic:

$$\lambda = \nu \, (\Phi_R^{\max} - \Phi_R). \tag{4}$$

where, $\nu$ characterizes the quality of the available nutrients and $\Phi_R^{max}$ is again the Y-offset, i.e. the maximal ribosome content that is theoretically possible.

Scott *et al.* propose that the simplest way to account for these linear growth laws is to assume a partition of the proteome in three fractions. The existence of a constant fraction $\Phi_Q$, that does not vary as the growth rate changes, is deduced from the fact that the ribosomal fraction cannot be increased further than a maximum of $\Phi_R^{\max} = 1 - \Phi_Q$. This fixed fraction is hypothesized to represent house-keeping proteins, whose concentration is kept constant by negative feedback mechanisms. The ribosomes and associated proteins required for translation constitute a further significant fraction of the proteome, $\Phi_R$. It's magnitude is closely associated to the growth rate of the cell, as described by the growth laws, equations 3 and 4. Finally, since $\Phi_Q$ is fixed, there needs to be a third fraction that accommodates for changes in $\Phi_R$. This variable fraction,

$$\Phi_P = 1 - \Phi_Q - \Phi_R = \Phi_R^{\max} - \Phi_R = \frac{\lambda}{\nu} \tag{5}$$

is interpreted as the group of proteins responsible for catabolism, as well as other proteins whose expression is unregulated, i.e. not subject to any of the homeostatic mechanisms that stabilize components of the Q-fraction. It follows the exact opposite behavior to the ribosomal fraction $\Phi_R$, i.e. decreasing with increasing nutrient quality and decreasing with increasing inhibition of ribosomes. Crucially for the derivation of our model framework below, constitutive genes were shown to belong to this category(2) (Figure S21).

The growth laws, equations 3 and 4, together with the constraint of having a finite proteome size,

$$\Phi_Q + \Phi_R + \Phi_P = 1 \tag{6}$$

are solved by Scott *et al.* to arrive at expressions for the growth rate and ribosomal fraction, that depend only on three parameters: the nutritional capacity $\nu$, the translational capacity $\gamma_0$ and the maximally-possible ribosomal fraction $\Phi_R^{\max}$.

$$\lambda \, (\gamma_0, \nu, \Phi_R^{\max}) = (\Phi_R^{\max} - \Phi_{R_0}) \, \frac{\gamma_0 \, \nu}{\gamma_0 + \nu} \tag{7}$$

$$\Phi_R \, (\gamma_0, \nu, \Phi_R^{\max}) = (\Phi_R^{\max} - \Phi_{R_0}) \, \frac{\nu}{\gamma_0 + \nu} + \Phi_{R_0} \tag{8}$$

Finally, Scott *et al.* also discussed what would happen if non-toxic, exogenous proteins are expressed in the cell. Exogenous gene expression would introduce a further proteome fraction, $\Phi_S$ ($S$ for *synthetic*), which results in growth defects because it restricts the fraction of the proteome that is available for ribosomes and catabolic proteins (effectively decreasing the value of $\Phi_R^{\max}$). With the extra proteome fraction, the finite proteome constraint becomes

$$\Phi_Q + \Phi_R + \Phi_P + \Phi_S = 1 \tag{9}$$

which leads to equations for the growth rate and ribosomal fraction that explicitly depend on $\Phi_S$:

$$\lambda\left(\gamma_0, \nu, \Phi_R^{\max}, \Phi_S\right) = \left(\Phi_R^{\max} - \Phi_{R_0} - \Phi_S\right) \frac{\gamma_0\, \nu}{\gamma_0 + \nu} \tag{10}$$

$$\Phi_R\left(\gamma_0, \nu, \Phi_R^{\max}, \Phi_S\right) = \left(\Phi_R^{\max} - \Phi_{R_0} - \Phi_S\right) \frac{\nu}{\gamma_0 + \nu} + \Phi_{R_0} \tag{11}$$

Note that this assumes a synthetic proteome fraction that is fixed, as is the case when a neutral protein product is expressed constitutively. Furthermore, $\Phi_S$ enters the equations as an extra parameter. Scott *et al.* express an exogenous protein in *E. coli* and measure the fraction of the proteome occupied by that protein. They show that the relation between that fraction and the growth defects observed can be quantitatively described by equation 10.

The parameters $\nu$, $\gamma_0$ and $\Phi_R^{\max}$ were estimated by Scott *et al.* for a wide range of conditions encompassing cells growing in different media and with different concentrations of chloramphenicol. Most studies on synthetic genetic circuits in *E. coli* are conducted in a laboratory setting under comparable conditions, e.g. the performance of a circuit is commonly evaluated in cells that grow exponentially at 37°C with saturated availability of oxygen and a well-defined growth medium. Under such conditions, $\gamma_0$ and $\Phi_R^{\max}$ do not vary much, which makes it possible to use the values estimated in (2). The above equations can then be used to predict growth defects due to burden, provided that the amount of synthetic proteins being expressed is known (with respect to the total protein mass of the cell) and the single missing parameter, $\nu$, is determined. $\nu$ can be estimated by measuring the growth rate of the strain of interest in the chosen media.

#### Host-aware modeling framework for synthetic genetic circuits

Expression of heterologous genes consumes cellular resources and reduces the growth rate of cells. In turn, limitations in the cellular gene expression machinery impact the performance of synthetic circuits(9). Here, we develop a modeling framework that captures this two-way interference and that can be seamlessly incorporated into conventional ODE models with little extra complexity and adding no extra free parameters.

In the proteome allocation model of Scott *et al.* summarized above, equations 10 and 11 quantify the impact of expressing heterologous genes on the cellular resources ($\Phi_R$) and the growth rate ($\lambda$). They are valid for balanced exponential growth. However, modelers are often interested in understanding the dynamics of circuits and not only their steady-state behavior. Moreover, the proteome fraction occupied by the circuit, $\Phi_S$, which enters the equations as a free parameter, cannot be determined experimentally in most practical cases. The equations also do not explicitly quantify the effect that the limited pool of cellular resources has on the performance of a circuit.

#### Derivation of the framework

In the following, we assume that the relations presented in the previous section also apply to the transient dynamics of a synthetic genetic circuit and not only at steady-state. To address the other issues, we begin by considering the simplest case of a circuit composed by a single, constitutively-expressed gene, $x$. The study by Scott *et al.* demonstrated that this type of unregulated expression would fall under the P-type proteome fraction(2, 7), which follows the opposite rules that apply to the ribosomal fraction (eq. 5). This means that an appropriate host-aware model of this simple circuit should recover two general trends:

1. the proteome fraction of $X$ should decrease linearly with growth rate, when growth is modulated by nutrient

quality

2. The proteome fraction of $X$ should increase linearly with growth rate, when growth is modulated by translational capacity and the slope of this increase should depend on the nutrient quality.

Conventional models of constitutive expression represent transcription and translation as birth-death processes:

$$\begin{aligned} \frac{dm_X}{dt} &= \omega - \delta_m \, m_X \\ \frac{dX}{dt} &= \alpha \, m_X - \lambda \, X \end{aligned} \tag{12}$$

where mRNA is transcribed at a rate $\omega$ and degraded at a rate $\delta_m$ and proteins are translated at a rate $\alpha$ and diluted through cell growth. At steady-state, the synthetic proteome fraction is given by

$$\Phi_S = \Phi_X = \frac{n_X}{\rho_{cell}} \, \overline{X} = \frac{n_X \, \omega \, \alpha}{\rho_{cell} \, \delta_m} \, \frac{1}{\lambda} \tag{13}$$

which clearly does not meet the requirements stated above (Figure S21A), since it scales as $1/\lambda$.

Our goal is to find a simple expression that captures how the required links between growth rate and constitutive gene expression listed above, may arise from coupling the circuit to the gene-expression machinery of the cell. As a first step, we introduce a dependency of the circuit on the cellular translation machinery, which is known to be the most important resource bottleneck(10) in *E. coli.* For simplicity, we assume mass action kinetics, with $R$ representing a unit of *active* translational resources[1] (active ribosome plus associated factors required for protein production). The ODEs become[2]

$$\begin{aligned} \frac{dm_X}{dt} &= \omega - \delta_m \, m_X \\ \frac{dX}{dt} &= \alpha \, m_X \, R - \lambda \, X \end{aligned} \tag{14}$$

We can convert the concentration of active translational machinery to its proteome mass fraction $(\Phi_R - \Phi_{R_0})$ and then use the first growth law (eq. 3) to express it as a function of the growth rate

$$R = \frac{\rho_{cell}}{n_R} (\Phi - \Phi_{R_0}) = \frac{\rho_{cell}}{n_R \, \gamma_0} \, \lambda \tag{15}$$

With this, the ODE for the protein concentration becomes

$$\frac{dX}{dt} = \alpha \, \frac{\rho_{cell}}{n_R \, \gamma_0} \, m_X \, \lambda \, - \lambda \, X \tag{16}$$

which would imply that at steady-state the concentration (and mass fraction) of the constitutive protein is independent of the growth rate (Figure S21B),

$$\Phi_S = \frac{n_X}{\rho_{cell}} \, \overline{X} = \frac{n_X \, \omega \, \alpha}{n_R \, \gamma_0 \, \delta_m} \tag{17}$$

[1] For a possible derivation of this mass-action term from a model of peptide elongation, see(11)

[2] To keep notation simple, we have left $\alpha$ as a symbol for the translation rate, although it must be noted that it now has different units compared to the quantity in eq. 12. The same will be true below, when we consider the role of transcriptional resources and the dependency on the media quality.

again not capturing the expected behavior of a constitutive gene.

In order to determine what the correct expression would look like, we take a more careful look at equation 13. The synthetic proteome fraction scales inversely with growth rate, because the dilution term is set by the growth rate. From equation 13, then we can infer that the production term in the ODE describing the protein concentration must scale as $\frac{\lambda^2}{\nu}$, if the synthetic proteome fraction is to be proportional to the P-type fraction

$$\Phi_S \propto \Phi_P = \frac{\lambda}{\nu} \tag{18}$$

We interpret this result in the following way. The squared dependency on growth rate arises quite naturally if we consider that the availability of transcriptional machinery also depends on the cellular growth rate. Here, we refer to a recent study by Balakrishnan *et al.*, in which the authors monitor transcriptome- and proteome-wide changes under different growth conditions, in order to investigate at a very detailed level how the observed growth-rate dependencies arise naturally from the central dogma of molecular biology(12).

Balakrishnan *et al.* report that the mRNA and protein levels correlate linearly for the majority of genes in *E. coli.* Moreover, after carefully quantifying the contributions of each step of gene expression, they conclude that the concentrations of proteins are dominantly set by the transcriptional output of their genes and not by other factors, such as translation initiation rates, mRNA degradation rates or gene dosage in replicating chromosomes. Balakrishnan *et al.* further estimate that the concentration of active RNA polymerase is tightly coupled to the concentration of active ribosomes. Therefore, it exhibits the same dependencies on growth rate as the ribosomal proteome fraction, i.e. the growth laws described in the previous section. Since the actual concentration of RNA polymerase components seems to remain constant under different growth conditions, the authors present evidence for a mechanism by which the availability of *active* RNA polymerase is modulated by the anti-sigma factor Rsd, which indeed exhibits the expected growth-rate dependencies.

Based on the findings of Balakrishnan *et al.*, we now consider that the transcription rate of the constitutive gene reflects the overall availability of RNA polymerase, as is the case for the translation rate and the active ribosomes

$$\frac{dm_X}{dt} = \omega \, \mathrm{RNAP} - \delta_m \, m_X \tag{19}$$

Since Balakrishnan *et al.* determined that the concentration of active RNA polymerase correlates positively with growth rate, we set

$$\mathrm{RNAP} = \beta \, \lambda \tag{20}$$

so that the ODE describing the mRNA concentration becomes

$$\frac{dm_X}{dt} = \widetilde{\omega} \, \lambda - \delta_m \, m_X \tag{21}$$

where we have absorbed the proportionality factor into the transcription rate.

With this, the equation for the synthetic proteome fraction becomes

$$\Phi_S = \frac{n_X}{\rho_{cell}} \, \overline{X} = \frac{n_X \, \widetilde{\omega} \, \alpha}{n_R \, \gamma_0 \, \delta_m} \, \lambda \tag{22}$$

This expression looks already more promising, since it is proportional to the growth rate. However, it does not distinguish between cases in which the growth rate is varied through changes in nutritional capacity or translational capacity (Figure S21C). Comparison to eq. 18 reveals that in order to achieve a behavior that does distinguish these two cases, there is still a factor $\frac{1}{\nu}$ missing.

It is less straightforward to derive the presence of this factor from first principles. However, we note that Scott *et al.* propose that the observed difference between the behavior of the P-sector and the R-sector reflects the balance between catabolic and anabolic fluxes in *E. coli*(2, 3). The factor $\frac{1}{\nu}$ captures this overall regulatory strategy, which affects a large portion of genes in parallel.

Since Balakrishnan *et al.* provide evidence that the concentrations of proteins are set foremost by the transcriptional activity, we here assume that the factor $\frac{1}{\nu}$ enters at that stage of the gene expression process. The final model for the expression of a constitutive gene thus becomes

$$\begin{aligned} \frac{dm_X}{dt} &= \widetilde{\omega}\,\frac{\lambda}{\nu} - \delta_m\, m_X \\ \frac{dX}{dt} &= \widetilde{\alpha}\, m_X\, \lambda - \lambda\, X \end{aligned} \tag{23}$$

where we have absorbed the constants $\frac{\rho_{cell}}{n_R\,\gamma_0}$ into an effective translation rate $\widetilde{\alpha}$.

These final equations have the same number of free parameters than the conventional model of the constitutive gene we started from. The nutritional capacity $\nu$ is a property of the media in which the cells are grown and it can be easily determined by measuring the growth rate of the strain in the absence of the circuit and then solving for $\nu$ in eq.10. We note that the choice of adding the factor $\frac{1}{\nu}$ to the transcriptional step is arbitrary. It could also be added to the translation step and still the predictions at the protein level would be consistent with the bacterial growth laws. Moreover, if the circuit of interest operates in a context where the media is fixed and there are no changes in nutrient quality, it becomes unnecessary to consider the factor $\frac{1}{\nu}$, since it can be absorbed into the production constants $\widetilde{\omega}$ or $\widetilde{\alpha}$.

Here, we have simply added an explicit growth-rate dependency to the production rates of mRNA and protein to reflect the fact that these depend on the physiological state of the cell through their reliance on the host's gene expression machinery. As Figure S21D shows, our modified equations are able to capture the expected qualitative behavior of a constitutive gene, both in the case when media quality is modulated and when the translational machinery is inhibited by a bacteriostatic antibiotic. Since the general dependence on the host gene-expression machinery does not only apply to constitutive genes, but to any heterologous gene that forms part of a synthetic network of genes[3], in the following selection we generalize the framework derived here to an arbitrary synthetic genetic circuit.

#### General Formulation of the framework

The host-aware framework we propose here can be applied to any arbitrary synthetic genetic circuit. It consist of simple modifications to a conventional ODE model, which incorporate a two-way coupling between the circuit and the host. The circuit impacts the host by sequestering cellular resources, which causes a reduction in the growth rate. The host influences the performance of the circuit, because the latter depends on the availability of cellular gene-expression machinery and this is tightly regulated in accordance to the host's physiological state.

Consider a generic synthetic genetic circuit composed of $u$ genes that are introduced into *E. coli* to produce proteins $X_1, \cdots, X_u$. A conventional model of such a system will consist of $2u$ ordinary differential equations, describing the mRNA and protein concentrations of the circuit's components. To these ODEs, we add the factors derived in the previous section to the production rates of mRNA and protein, so that they take the form

[3]This excludes genes whose expression does not depend on cellular factors, such as genes transcribed by exogenous polymerases or translated by orthogonal ribosomes(13).

$$\begin{aligned}\frac{dm_{X_i}}{dt} &= \widetilde{\omega}_i\, T_i(\boldsymbol{m_X}, \boldsymbol{X})\, \frac{\lambda}{\nu} + F_i(\boldsymbol{m_X}, \boldsymbol{X}) - \delta_i\, m_{X_i} \\ \frac{dX_i}{dt} &= \widetilde{\alpha}_i\, m_{X_i}\, \lambda + G_i(\boldsymbol{m_X}, \boldsymbol{X}) - \lambda\, X_i\end{aligned} \tag{24}$$

with $i = 1, \cdots u$. The collections of functions $\boldsymbol{F}(\boldsymbol{m_X}, \boldsymbol{X})$ and $\boldsymbol{G}(\boldsymbol{m_X}, \boldsymbol{X})$ describe potential interactions between the components of the circuits at the mRNA and protein level, respectively, and the functions $\boldsymbol{T}(\boldsymbol{m_X}, \boldsymbol{X})$ reflect potential regulatory input at the transcriptional level.

The explicit dependency of the production and dilution rates on the growth rate reflects the impact of resource availability and host growth on the concentration of circuit components. To get an explicit expression for the growth rate, which reflects the impact of the circuit on the host, we make use of the expressions derived by Scott *et al.*[4] for the case of heterologous gene expression (eq. 10).

We can use the conversion factor derived above to calculate the proteome fraction occupied by the proteins in the circuit at any particular point in time:

$$\Phi_S = \frac{\sum_{i=1}^{u} n_{X_i}\, X_i}{\rho_{cell}} \tag{25}$$

This can in turn be inserted into equation 10 to obtain an explicit expression for the instantaneous growth rate

$$\lambda\,(\Phi_S) = (\Phi_R^{\max} - \Phi_{R_0} - \Phi_S)\, \frac{\gamma_0\, \nu}{\gamma_0 + \nu} \tag{26}$$

Equations 24, 25 and 26 form a closed system of $2u$ ODEs and 2 algebraic equations that can be simulated to approximate the dynamic behavior of the synthetic genetic circuit in the context of an exponentially growing *E. coli* host.

### Model parameters

Under the growth conditions investigated by Scott *et al.*(2), the estimated values for $\Phi_R^{\max}$, $\Phi_{R_0}$ and $\gamma_0$ were largely constant. As long as the cells under consideration are grown under similar conditions[5], which will be true of most cases in a laboratory environment, the values they determined for these three parameters can be used.

Apart from these, the number of amino acids that compose each protein, $n_{X_i}$ is known, as well as the cellular protein density $\rho_{cell}$, which is largely growth-rate independent. The nutritional capacity $\nu$, which describes the quality of the media, can be easily determined by measuring the growth rate of the strain in the absence of the synthetic genetic circuit ($\Phi_S = 0$) and substituting in equation 26:

$$\nu = \frac{\gamma_0\, \lambda}{\gamma_0\, (\Phi_R^{\max} - \Phi_{R_0}) - \lambda} \tag{27}$$

In that case, the framework we propose contains the same amount of free parameters as a conventional model that does not consider the host, that is $u$ transcription rates, $u$ mRNA degradation rates, $u$ translation rates plus any number of parameters required to describe the interactions between circuit species. This framework then has the advantage that it can be easily incorporated into an existing ODE model and that it can capture the qualitative trends

[4] This assumes that the dynamics of the synthetic circuit can be taken to be in quasi-equilibrium compared to the dynamics of the host's physiology. We simply assume this here, since this results in model predictions that match the experimental data for our circuit of interest.

[5] Carbon-limited exponential growth at 37°C with saturating amounts of carbon source, in the presence or absence of translation-inhibiting antibiotics.

expected from a host-aware model without adding complexity or increasing the risk of overfitting. The parameter values we use in this study are summarized in table 1.

#### Model of photophilic strain

In this section, we apply the framework formulated above to the case of the growth-control circuit in the photophilic strain. The circuit consists of the two parts of a split opto-T7, which dimerizes in the presence of blue light to produce an active complex that transcribes mRNA coding for the resistance enzyme chloramphenicol acetyltransferase (CAT).

We begin by presenting the final result and then derive the equations in the following sections. Our model is composed of three ODEs and two algebraic equations:

$$\frac{dT_T}{dt} = \hat{\alpha}_T \, \frac{\lambda^2}{\nu} - \lambda \, T_T \tag{28}$$

$$\frac{dT_D}{dt} = h_{ON}(L) \, \left(T_T - 2 \, (T_D + g_{ON}(T_D))\right)^2 - \lambda \, (T_D + g_{ON}(T_D)) \tag{29}$$

$$\frac{dC}{dt} = \hat{\alpha}_C \, \frac{\lambda}{\nu} \, g_{ON}(T_D) - \lambda \, C \tag{30}$$

$$\Phi_S(t) = \frac{n_T \, T_T(t) + n_C \, C(t)}{\rho_{cell}} \tag{31}$$

$$\lambda(t) = (\Phi_R^{max} - \Phi_{R_0} - \Phi_S(t)) \, \frac{\nu \, \gamma_0}{\gamma_0 + \nu \left(1 + \frac{\frac{A_E}{K_D}}{1 + \left(\frac{C(t)}{\kappa \, \hat{K}_C}\right)^{h_C}}\right)} \tag{32}$$

The ODEs describe the evolution of the concentration of the circuit's proteins: T7-monomers, $T_T$, T7-dimers, $T_D$ and CAT, $C$. For simplicity, we have assumed fast mRNA dynamics for all species (quasi-steady state), which in our framework corresponds to:

$$m_{X_i}^{qss} = \frac{\widetilde{\omega}_i}{\delta_i} \, \frac{\lambda}{\nu} \tag{33}$$

and we have absorbed the transcription and mRNA-degradation rates into an effective protein-production rate $\widetilde{\alpha}_i$.

The light-dependent T7-dimerization rate is modeled by the general expression

$$h_{ON}(L) = h_{ON}^{min} + \left(h_{ON}^{max} - h_{ON}^{min}\right) \, \frac{L^{n_L}}{K_L^{n_L} + L^{n_L}} \tag{34}$$

where $L$ is the blue-light intensity, and the factor

$$g_{ON}(T_D) = N_p \, \frac{T_D^{n_G}}{K_G^{n_G} + T_D^{n_G}} \tag{35}$$

captures the concentration of actively transcribing T7-dimers.

The difference between equation 32 and equation 26 reflects the effect of chloramphenicol inhibition.

The equations used here are mainly phenomenological in nature. Nevertheless, their utility is supported by the good results obtained, both in recapitulating the gene expression and growth rate dynamics of the photophilic strain, as well as in predicting the trajectories of the closed-loop co-culture. In the following sections, we provide the rationale behind the choice of terms for our model.

#### Production of CAT

To get a simple description of transcription by T7 polymerase, we treat it as a two step process

$$T_D + g_{OFF} \underset{\beta_-}{\overset{\beta_+}{\rightleftharpoons}} g_{ON} \xrightarrow{\widetilde{\omega}_C} T_D + g_{OFF} + m_C$$

First, a T7-dimer first binds reversibly to a copy of the gene which is in an inactive state, $g_{OFF}$, to produce an active complex, $g_{ON}$. The complex then directly produces a unit of CAT mRNA, $m_C$, thereby releasing the T7-dimer and the gene, which reverts back to its OFF state. Finally, the CAT gene resides on a plasmid, whose copy number, $N_P = g_{OFF} + g_{ON}$, we assume is kept constant through a balance between plasmid replication and dilution:

$$\varnothing \xrightarrow{\lambda\, N_p} g_{OFF} \xrightarrow{\lambda} \varnothing$$
$$g_{ON} \xrightarrow{\lambda} \varnothing$$

These reactions lead to the following ODEs for the time evolution of the inactive and active gene copies

$$\begin{aligned} \frac{dg_{OFF}}{dt} &= \lambda\, N_p - \beta_+\, g_{OFF}\, T_D + (\beta_- + \widetilde{\omega}_C)\, g_{ON} - \lambda\, g_{OFF} \\ \frac{dg_{ON}}{dt} &= \beta_+\, g_{OFF}\, T_D - (\beta_- + \widetilde{\omega}_C)\, g_{ON} - \lambda\, g_{ON} \end{aligned} \tag{36}$$

Assuming that these dynamics evolve at a faster timescale than that of processes related to the changes in cell growth and physiology, we consider steady-state for these equations, which leads to the following expression for the concentration of active gene copies

$$g_{ON} = N_p\, \frac{T_D}{T_D + \frac{\beta_- + \widetilde{\omega}_C}{\beta_+} - \frac{\lambda}{\beta_+}} \approx N_p\, \frac{T_D}{T_D + K_G} \tag{37}$$

where we have used the fact that the binding reaction of the T7 to its promoter is orders of magnitude faster than cell growth and we have introduced $K_G = \frac{\beta_- + \widetilde{\omega}_C}{\beta_+}$ as a parameter that relates to the affinity of the T7 to its promoter sequence. We finally add a hill-coefficient to this expression to be able to capture higher degrees of non-linearity in the system, leading to the expression in equation 35.

With this, and assuming the mRNA concentration is in quasi-steady state (eq. 33), we can write the ODE for the concentration of CAT as

$$\frac{dC}{dt} = \hat{\alpha}_C\, \frac{\lambda}{\nu}\, N_p\, \frac{T_D^{n_G}}{T_D^{n_G} + K_G^{n_G}} - \lambda\, C \tag{38}$$

where we have again absorbed the parameters related to transcription into the effective production rate $\hat{\alpha}_C$. Note that, in this equation, the protein production rate scales only linearly with the growth rate. This is because, in the derivation of our framework, the square dependency arises from the fact that both the ribosomes and RNA polymerase introduce a linear dependency on the growth rate. However, since CAT is not transcribed by the endogenous *E. coli* polymerase, we let it scale linearly with the growth rate to reflect the fact that it only depends on the cellular translation machinery.

#### Light-dependent dimerization

We begin by representing light-mediated dimerization of the split T7 through the following reversible reaction

$$T_M + T_M \underset{h_{OFF}(L)}{\overset{h_{ON}(L)}{\rightleftharpoons}} T_D$$

In reality, the opto-T7 is a heterodimer, composed of two different N-terminal, nMag-T7$_{\text{N-Ter}}$, and C-terminal, pMag-T7$_{\text{C-Ter}}$, parts. However, for simplicity we assume that we can model the opto-T7 as a homodimer that results from the light-promoted fusion of two identical monomer units, $T_M$.

We chose a general Hill-function, each with 4 free parameters, for the forward and backward rates of the dimerization reaction, in order to capture the dynamics of this process accurately

$$\begin{aligned} h_{ON}(L) &= h_{ON}^{min} + \left(h_{ON}^{max} - h_{ON}^{min}\right) \frac{L^{n_{L_{ON}}}}{K_L^{n_{L_{ON}}} + L^{n_{L_{ON}}}} \\ h_{OFF}(L) &= h_{OFF}^{max} - \left(h_{OFF}^{max} - h_{OFF}^{min}\right) \frac{L^{n_{L_{OFF}}}}{K_L^{n_{L_{OFF}}} + L^{n_{L_{OFF}}}} \end{aligned} \tag{39}$$

The ODE for the T7 dimers includes the dimerization and transcription reactions

$$\frac{dT_D}{dt} = h_{ON}(L)\,T_M^2 - h_{OFF}(L)\,T_D - \beta_+\,g_{OFF}\,T_D + (\beta_- + \widetilde{\omega}_C)\,g_{ON} - \lambda\,T_D \tag{40}$$

$$= h_{ON}(L)\,T_M^2 - h_{OFF}(L)\,T_D - \lambda\,g_{ON} - \lambda\,T_D \tag{41}$$

However, simulations revealed that the dynamics of $T_D$ were only weakly influenced by the dissociation reaction, in contrast to the forward reaction, which scales with the square of the monomer concentration. Because of this and in order to reduce the number of free parameters, we neglected the dissociation reaction, effectively considering dimerization as an irreversible process. This leads to the simpler equation

$$\frac{dT_D}{dt} = h_{ON}(L)\,T_M^2 - \lambda\,(g_{ON} + T_D) \tag{42}$$

Finally, we chose to keep track of the concentration of *total* T7-monomers, $T_T$, instead of that of *free* T7-monomers, $T_M$. $T_T$ is given by

$$T_T = T_M + 2\,(T_D + g_{ON}) \tag{43}$$

and its concentration is not affected by dimerization or transcription reactions, so that its dynamics follow the simple ODE

$$\frac{dT_T}{dt} = \hat{\alpha}_T\,\frac{\lambda^2}{\nu} - \lambda\,T_T \tag{44}$$

which is equation 28. Using equation 43 to remove $T_M$ from the ODE for $T_D$, we recover equation 29.

#### Inhibition of growth by chloramphenicol

What remains to be discussed is how to model the inhibitory effect of chloramphenicol on growth and how this is alleviated in the presence of the resistance. For this, we follow closely the approach taken by Deris *et al.*(7) and we refer to their paper for further details.

Chloramphenicol inhibits translational elongation by direct binding to the 50S subunit of the ribosome. We model this by considering that the ribosomes can be in one of three states. $R_u$ are active, chloramphenicol-unbound ribosomes which can be engaged in translation. $R_b$ are ribosomes that have been inhibited by antibiotic binding and $R_0$ are the naturally-occurring inactive ribosomes that give rise to the Y-offset in the bacterial growth law, eq. 3. We model chloramphenicol inhibition as a reversible binding reaction

$$R_u + A_I \underset{k_-}{\overset{k_+}{\rightleftharpoons}} R_b \xrightarrow{\lambda} \varnothing$$

where $A_I$ is the intracellular chloramphenicol concentration. This leads to the following ODE for the concentration of the bound ribosomal species

$$\frac{dR_b}{dt} = k_+ \, R_u \, A_I - k_- \, R_b - \lambda \, R_b \tag{45}$$

Reversible chloramphenicol binding happens at a faster timescale than growth and gene expression. Therefore, we assume $R_b$ to be in quasi-steady state, from which we get

$$R_u = \frac{\left(1 + \frac{\lambda}{k_-}\right) \, (R - R_0)}{1 + \frac{k_+}{k_-} \, A_I + \frac{\lambda}{k_-}} \approx \frac{1}{1 + \frac{A_I}{K_D}} \, (R - R_0) \tag{46}$$

where we have used the fact that $k_- \gg \lambda$ and we have introduced the total ribosomal concentration $R = R_u + R_b + R_0$ and the parameter $K_D = \frac{k_-}{k_+}$, which relates to the affinity of chloramphenicol to the ribosome. Equation 46 reveals that the presence of chloramphenicol reduces the concentration of actively translating ribosomes by a factor $\frac{1}{1+\frac{A_I}{K_D}}$.

Since we can transform ribosomal concentrations to proteome mass fractions by multiplication with the factor $\frac{n_R}{\rho_{cell}}$, the same applies to the corresponding mass fractions

$$\Phi_{R_u} = \frac{1}{1 + \frac{A_I}{K_D}} \, (\Phi_R - \Phi_{R_0}) \tag{47}$$

In the presence of chloramphenicol, only unbound ribosomes contribute to biomass synthesis, so the first bacterial growth law becomes

$$\lambda = \gamma_0 \, \Phi_{R_u} = \frac{\gamma_0}{1 + \frac{A_I}{K_D}} \, (\Phi_R - \Phi_{R_0}) \tag{48}$$

Again, we combine this equation with the second growth law

$$\Phi_P = \frac{\lambda}{\nu}$$

and the finite size of the proteome

$$\Phi_Q + \Phi_R + \Phi_P + \Phi_S = 1$$

to obtain an updated equation for the instantaneous growth rate in the presence of chloramphenicol

$$\lambda = (\Phi_R^{max} - \Phi_{R_0} - \Phi_S(t)) \, \frac{\nu \, \gamma_0}{\gamma_0 + \nu \left(1 + \frac{A_I}{K_D}\right)} \tag{49}$$

Finally, we turn to how the intracellular concentration of chloramphenicol, $A_I$, can be related to the extracellular concentration, $A_E$, which is the one we actually set in experiments. Apart from the reversible binding to the ribosomes, we assume that chloramphenicol enters the cell passively via diffusion and is also degraded catalytically by the action of CAT

$$\varnothing \underset{\kappa}{\overset{\kappa\, A_E}{\rightleftharpoons}} A_I$$
$$A_I + C \underset{v_-}{\overset{v_+}{\rightleftharpoons}} A^* \xrightarrow{v_f} C$$

The ODEs that describe the collection of reactions involving chloramphenicol are the following

$$\begin{aligned} \frac{dA_I}{dt} &= \kappa(A_E - A_I) - v_+\, A_I\, C + v_-\, A^* - k_+\, A_I\, R_u + k_-\, R_b - \lambda\, A_I \\ \frac{dA^*}{dt} &= v_+\, A_I\, C - v_-\, A^* - v_f\, A^* - \lambda\, A^* \end{aligned} \tag{50}$$

Assuming that degradation is a fast process, we assume quasi-steady state for $A*$, which leads to

$$A^* = \frac{\frac{v_+}{v_- + v_f}}{1 + \frac{\lambda}{v_- + v_f}}\, A_I\, C \approx \frac{A_I\, C}{K_C} \tag{51}$$

where we have again used the fact that binding and degradation reactions are fast compared to cell growth, $v_- + v_f \gg \lambda$, and we have introduced the parameter $K_C = \frac{v_- + v_f}{v_+}$, which relates to the affinity of the degradation reaction.

Using this result, together with the quasi-steady state concentration of bound ribosomes from eq. 45, we obtain

$$\begin{aligned} \frac{dA_I}{dt} &= \kappa(A_E - A_I) - \frac{1 + \frac{\lambda}{v_f}}{\frac{K_C}{v_f}}\, A_I\, C - \lambda\, \frac{A_I\, R_u}{K_D} - \lambda\, A_I \\ &\approx \kappa\, A_E - A_I \left[\kappa + \frac{C}{\hat{K}_C} + \frac{\rho_{cell}}{n_r\, \gamma_0\, K_D}\, \lambda^2 + \lambda\right] \end{aligned} \tag{52}$$

where we use the first growth law to replace $R_u$ with $\frac{\rho_{cell}\, \lambda}{n_r\, \gamma_0}$, we absorb $v_f$ into $\hat{K}_C = \frac{v_- + v_f}{v_+\, v_f}$ and for the last approximation we use the fact that $v_f \gg \lambda$.

We now examine the value ranges each term in this equation can take, to determine the ones that dominate. For the sources from which we obtain our parameter values, see table 1.

- For the concentrations of chloramphenicol and light intensities used in our experiments, we obtain growth rate values in the range $\lambda \in [0.015,\, 0.033]\frac{1}{min}$
- With this, the term with square-dependency on the growth rates takes on values in the interval $[0.1382,\, 3]\frac{1}{min}$
- The diffusion rate $\kappa \approx 90\frac{1}{min}$ is one to two orders of magnitude larger than the last two terms
- Since $\hat{K}_C \approx 0.333\, nM\, min$, the term $\frac{C}{\hat{K}_C}$ scales as $3\, C$, which can become comparable to $\kappa$ or dominant for even low concentrations of CAT.

Because of this, we further approximate the ODE for $A_I$ as

$$\frac{dA_I}{dt} \approx \kappa\, A_E - \kappa\, A_I + \frac{C}{\hat{K}_C}\, A_I \tag{53}$$

which, since $A_I$ only takes part in fast binding or degradation reactions can be taken to be at quasi-steady state, leading to

$$A_I \approx \frac{A_E}{1 + \frac{C}{\kappa \hat{K}_C}} \tag{54}$$

Inserting equation 54 back into equation 49, we recover the expression for growth rate in our model, eq. 32, except for the exponent $h_C$, which we added *a posteriori* because it improved the capability of the model to capture the non-linear response of the photophilic strain in experiments.

#### Modeling the dynamics of co-culture composition

We consider an exponentially growing two-strain co-culture in a turbidostat setup. The accumulation of total biomass $M_T$ in the vial is given by the combined exponential growth of the two strains that are present (in this section, $p$ and $c$ subscripts denote the photophilic and constitutive strains respectively):

$$\frac{dM_T}{dt} = \lambda_p \, M_p + \lambda_c \, M_c$$

The principle of a turbidostat consists in balancing the biomass increase rate by an equal dilution rate, so that the overall density of the culture is kept constant. To get a formula for this effective rate of dilution, $d$, we set

$$\frac{dM_T}{dt} \stackrel{!}{=} d \, M_T$$

which leads to

$$d = \frac{\lambda_p \, M_p + \lambda_c \, M_c}{M_p + M_c} = \lambda_p \, \varphi_p + \lambda_c \, \varphi_c$$

where we have defined the strain fractions in the co-culture $\varphi_p = \frac{M_p}{M_p + M_c}$ and $\varphi_c = \frac{M_c}{M_p + M_c}$, with $\varphi_c + \varphi_p = 1$.

The differential equation governing the dynamics of the photophilic fraction, $\varphi_p$ can be derived as follows:

$$\frac{d\varphi_p}{dt} = \frac{d}{dt}\frac{M_p}{M_T} = \frac{1}{M_T}\frac{dM_p}{dt} + M_p \frac{d}{dt}\frac{1}{M_T} = \frac{1}{M_T}\frac{dM_p}{dt} = \frac{\lambda_p \, M_p - d \, M_p}{M_T} = (\lambda_p - d) \, \varphi_p$$

where we have used the fact that in turbidostat mode $\frac{dM_T}{dt} = 0$. Inserting the expression for the dilution rate $d$ from above and noting that $\varphi_c = 1 - \varphi_p$ we get the following ODE for $\varphi_p$:

$$\frac{d\varphi_p}{dt} = (\lambda_p - \lambda_c) \, (1 - \varphi_p) \, \varphi_p \tag{55}$$

The equation provides immediate intuition about the co-culture dynamics, with three qualitatively different scenarios which are schematically shown in Figure 5A of the main text. Dynamic equilibria arise when the derivative in equation 55 vanishes. This happens for $\varphi_p = 1$ and $\varphi_p = 0$, corresponding to the extinction of either the constitutive or the photophilic strain.

The stability of the fixed points is determined by the sign of $\frac{d\varphi_p}{dt}$ close to the fixed point. Since $0 \leq \varphi_p$ and $0 \leq 1 - \varphi_p$, the sign of the derivative is completely determined by the sign of the first factor. If $\lambda_c > \lambda_p$, the fixed point at $\varphi_p = 1$ becomes unstable and the co-culture evolves towards $\varphi_p = 0$, a culture dominated by the constitutive strain. Alternatively, if $\lambda_p > \lambda_c$, the fixed point at $\varphi_p = 0$ becomes unstable and the photophilic strain will inevitably come to dominate. In both of these cases, the magnitude of the difference in growth rates sets how fast the system converges to the stable equilibrium.

When both strains grow at the same pace ($\lambda_c = \lambda_p$), the derivative vanishes irrespective of the value of $\varphi_p$. This means that the co-culture will maintain any particular ratio that it has when $\lambda_c$ and $\lambda_p$ first become equal.

In view of this, the control strategy for stabilizing the strain ratio with optogenetic feedback becomes clear. If the ratio is lower than the desired value, light levels that result in the photophilic strain growing faster than the constitutive one should be applied, steering the system towards $\varphi = 1$. The opposite is true if the ratio is higher than the desired value. Then, once the desired setpoint is reached, the light level should be such as to precisely ensure that the growth rates are equal, so that the desired ratio is maintained over time.

The analysis also reveals the fragility of the co-existence equilibrium. Even small fluctuations in the growth rate will cause the system to evolve towards a monoculture.

For our simulations, $\lambda_c$ is assumed to be constant. On the contrary, $\lambda_p$ is calculated at any given time by solving the set of equations 29-32 for the current external light input $L$, in order to compute the instantaneous growth rate of the photophilic strain via equation 32.

### 2 Construction of *evotron* framework

#### 2.1 eVOLVER sleeve re-design

##### Hardware

We re-designed the eVOLVER sleeve as shown in Figure S3 and Figure S4 to accommodate new tube holder and vial cap designs resulting in stable and consistent OD sensor measurements (Figure S6). We also integrated one blue LED (465nm, superbrightleds: RL5-B08-360) per sleeve to facilitate optogenetic stimulation of target cell culture when placed in the sleeve. This blue LED was connected to the vial electronic board by replacing the 90 degree photodiode (IR detector), shown in Figure S4B and Figure S5A. Since, our experiments involved cell cultures maintained at lower cell densities (OD $< 0.2$), readings from 135 degree photodiode (IR detector) were enough to provide accurate OD measurements (thus, 90 degree photodiode was not required). Further, we also replaced the default M6 plugin (on the eVOLVER motherboard) with a PWM board(1) to power this additional LED, and re-programmed UC2 arduino (SAMD21) to provide software control access for changing LED intensity in real-time during optogenetic experiments.

##### Software

All CAD design files, codes and routines are available in this GitHub repository:
`https://github.com/santkumar/evotron.git`

#### 2.2 Automated sampling platform design

##### Hardware

We designed our automated sampling and measurement framework around Opentrons OT-2 liquid handling robot. The modified eVOLVER platform was placed inside the Opentron OT-2 robot replacing its deck, as can be seen in Figure S8A. Three cleaning solution bottles (1-$H_2O$, 2-Bleach (2%), 3-$H_2O$) were also placed on the OT-2 deck for cleaning the sample path after every sampling step. A sampling-needle was fixed onto the OT-2 pipette head by using a custom-designed 3D printed needle holder (Figure S8A).

As seen in Figure S8B, the sampling needle was attached to a flexible silicone tubing which provided a route for the sample to move into the flow-cytometer (Cytoflex S, Beckman Coulter) sampling vial via a peristaltic sampling pump (Intellicyt peristaltic pump). Another tubing and waste-removal pump was set up to remove the residual sample from flow-cytometer sampling vial after measurement. A controller (Arduino Mega 2560) was incorporated in the framework providing software control access of the OT-2 robot (serial communication), sampling and waste-removal pumps (TTL signaling). The control computer was connected to the OT-2 and eVOLVER platforms via WiFi communication, and was set up to communicate with the arduino controller serially (Figure 3B).

#### Software

All CAD design files, codes and routines are available in this GitHub repository:
`https://github.com/santkumar/evotron.git`

#### Pseudo Code

These are the synchronized steps performed during an automated sampling event:

1. Start the sampling pump.
2. Move the OT-2 pipette head to a desired eVOLVER sleeve location (where our target cell culture is maintained within OD range 0.1-0.15 in a glass vial).
3. Lower the OT-2 pipette head so that the sampling-needle is dipped into the cell culture, wait for 3 seconds (for the sampling pump to extract ∼0.5ml of culture into the sampling tubing), and then move it up.
4. Wait for 40 seconds so that the extracted sample is pulled and completely discharged into the flow-cytometer sampling vial, then stop the sampling pump.
5. Start the flow-cytometer measurement.
6. Once measurement is done, start the waste-removal pump and wait for 10 seconds so that the residual sample is completely pumped out of the flow-cytometer samling vial, and then stop the waste-removal pump.
7. Start the sampling pump.
8. Move the OT-2 pipette head to the first cleaning solution (sterile $H_2O$) bottle location.
9. Lower the OT-2 pipette head so that the sampling-needle is dipped into the cleaning solution, wait for 4 seconds (for the sampling pump to extract ∼0.7ml of cleaning solution into the sampling tubing), and then move it up.
10. Wait for 40 seconds so that the extracted cleaning solution is pulled and completely discharged into the flow-cytometer sampling vial, then stop the sampling pump.
11. Start the waste-removal pump and wait for 10 seconds so that the cleaning solution is completely pumped out of the flow-cytometer samling vial, and then stop the waste-removal pump.
12. Move the OT-2 pipette head to the second cleaning solution (2% bleach solution) bottle location.
13. Lower the OT-2 pipette head so that the sampling-needle is dipped into the bleach solution, wait for 6 seconds (for the sampling pump to extract ∼1ml of bleach solution into the sampling tubing), and then move it up.
14. Wait for 40 seconds so that the extracted bleach solution is pulled and completely discharged into the flow-cytometer sampling vial, then stop the sampling pump.
15. Wait for 10 seconds so that the bleach solution has sufficient time to disinfect the sampling tubing and the flow-cytometer sampling vial.

16. Start the waste-removal pump and wait for 10 seconds so that the bleach solution is completely pumped out of the flow-cytometer samling vial, and then stop the waste-removal pump.

17. Move the OT-2 pipette head to the third cleaning solution (sterile $H_2O$) bottle location.

18. Lower the OT-2 pipette head so that the sampling-needle is dipped into the cleaning solution, wait for 8 seconds (for the sampling pump to extract a bit more than 1ml of cleaning solution into the sampling tubing), and then move it up.

19. Wait for 40 seconds so that the extracted cleaning solution is pulled and completely discharged into the flow-cytometer sampling vial, then stop the sampling pump.

20. Start the waste-removal pump and wait for 10 seconds so that the cleaning solution is completely pumped out of the flow-cytometer samling vial, and then stop the waste-removal pump.

21. The automated sampler is now ready to perform the next sampling.

**Note:** During a closed-loop control experiment, the feedback controller computation routine is run on measurements just after the flow-cytometer measurement is done in step 6. Once the computation is over, the LED (integrated on the respective eVOLVER sleeve of the target cell culture vial) stimulation intensity is set as per the value computed by this controller. These controller computations are carried out in parallel with cleaning steps 7-20.

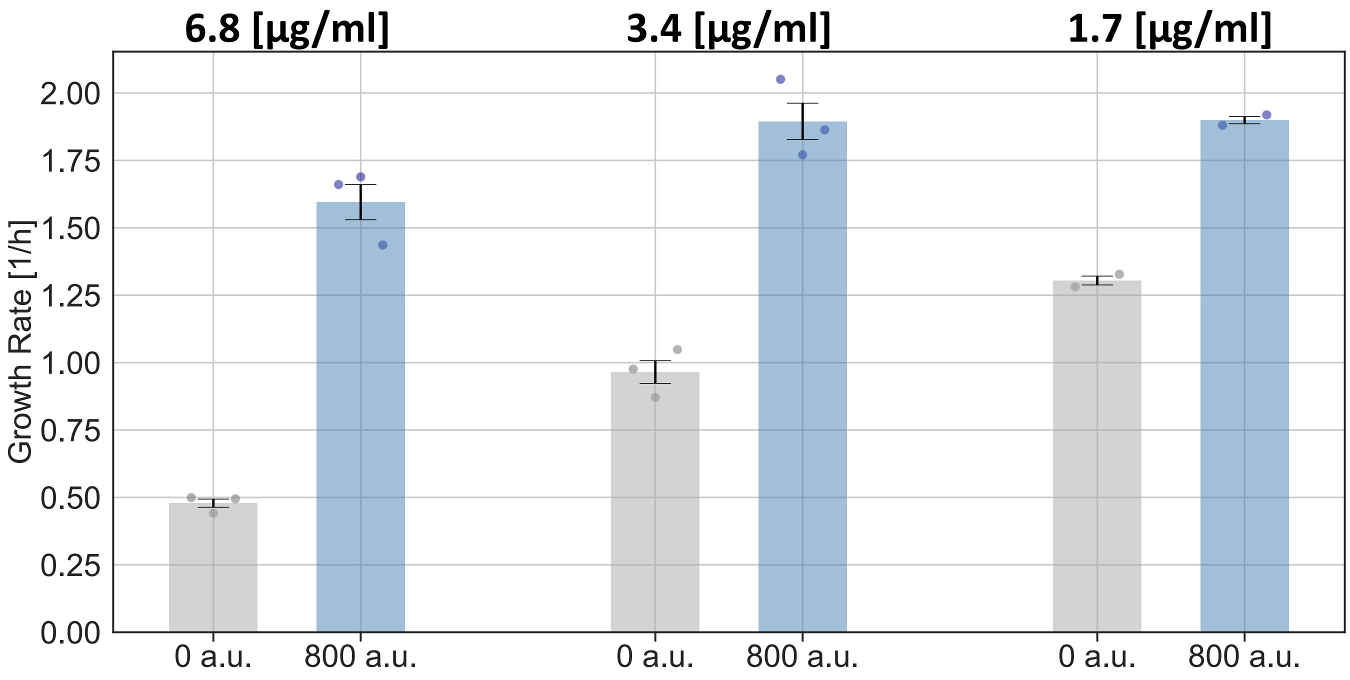


**Supplementary Fig. 1. Growth of photophilic strain with different external concentrations of chloramphenicol**
The growth rate of the photophilic strain can be modulated by changing the light intensity or the external concentration of chloramphenicol. Different bars correspond to equilibrium growth rate of the photophilic strain in the presence of different chloramphenicol concentrations and either no light or the maximal intensity used in this study (n=3 for 6.8 and 3.4 [ug/ml] and n=2 for 1.7 [ug/ml]). For the rest of the study, we used 3.4 ug/ml as fixed chloramphenicol concentration, because it leads to a good fold-change in growth, while maintaining a saturated growth rate when the resistance is fully induced. We hypothesized that this would reduce the selection pressure during closed-loop co-culture experiments for mutations that escape the growth control circuit.

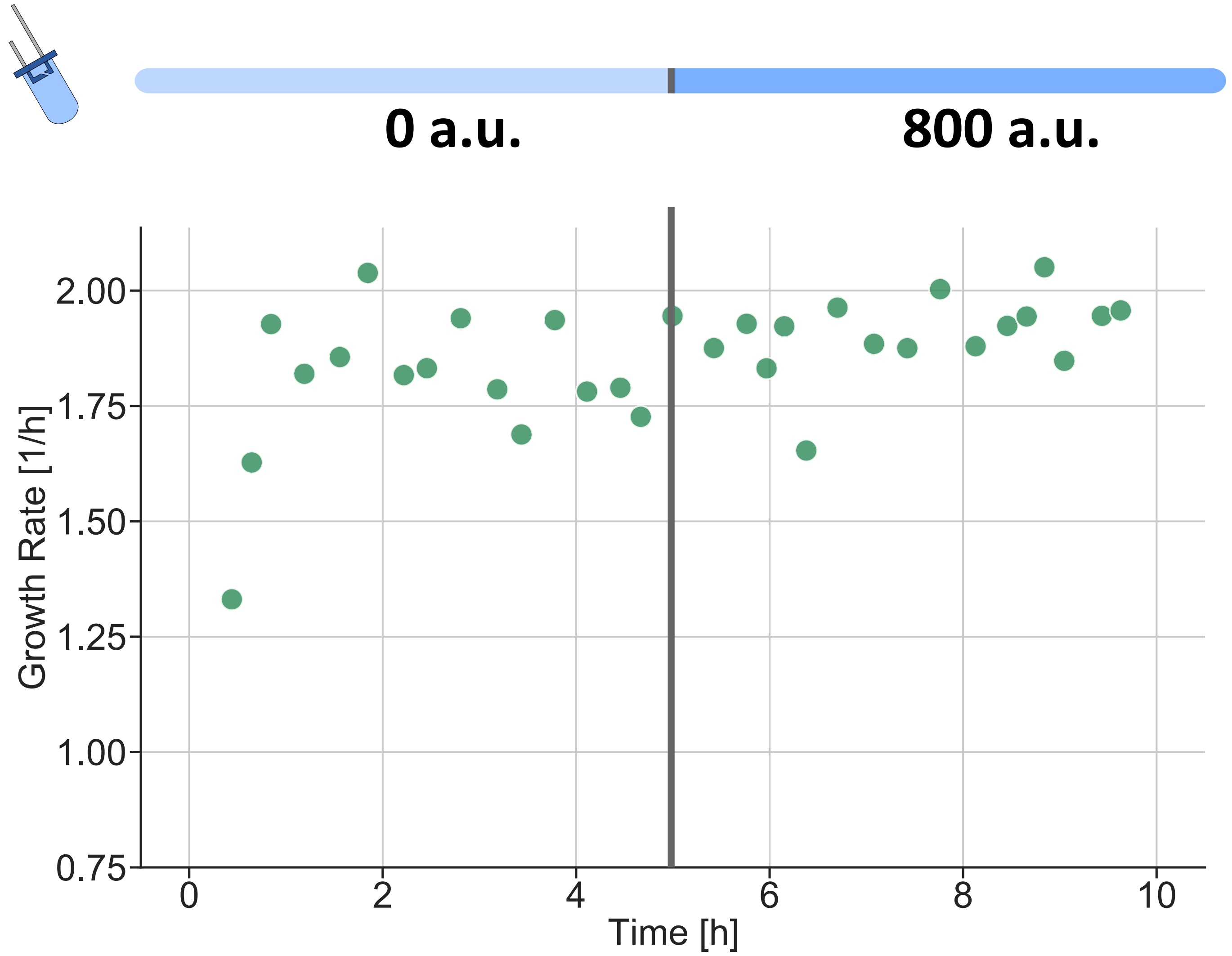


**Supplementary Fig. 2. Growth of photophilic strain is not affected by light in the absence of chloramphenicol**
The photophilic strain was grown in the absence of chloramphenicol and the culture was grown in the dark for 5h and then illuminated with maximum light intensity. Light does not cause any appreciable difference in the growth rate, suggesting that the induction of the resistance causes negligible metabolic burden and that there is no direct phototoxicity.

A

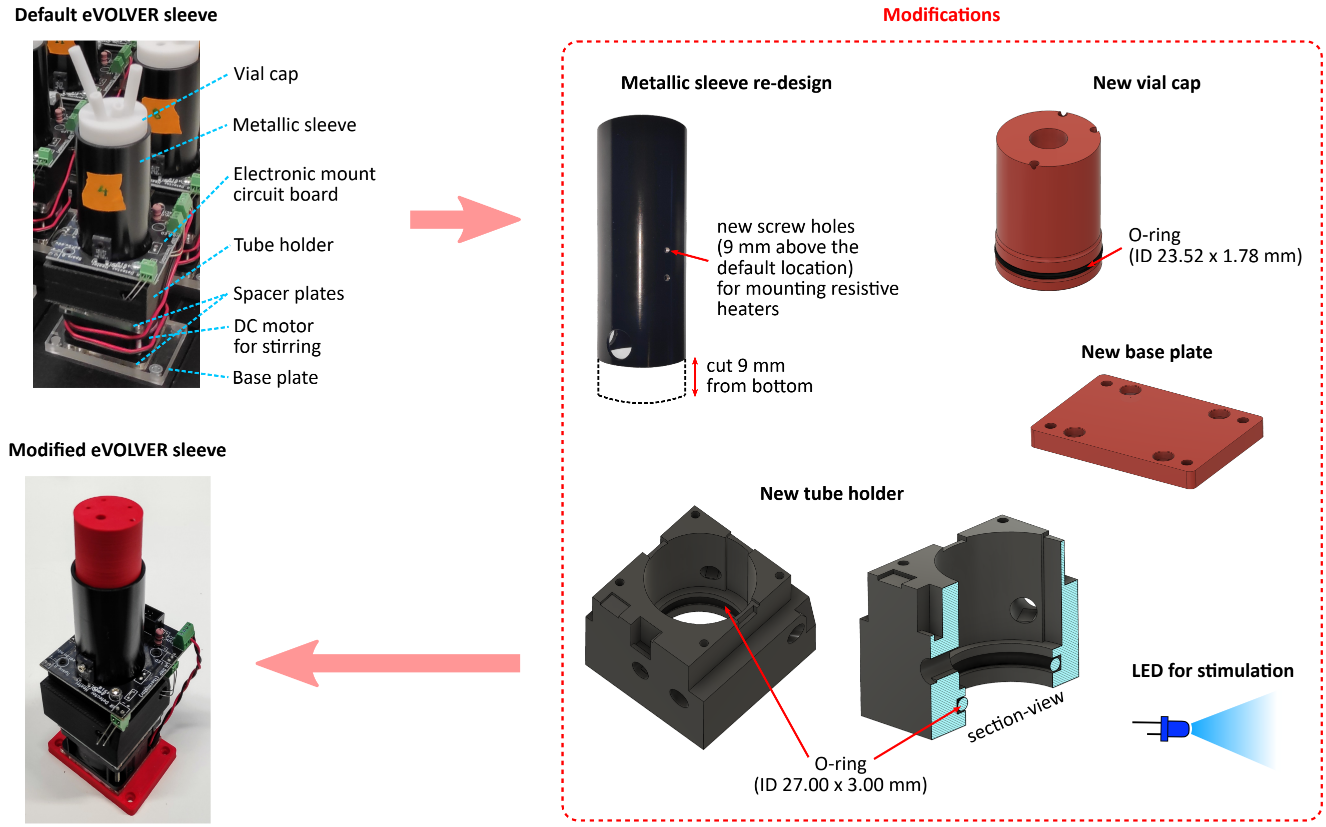


B

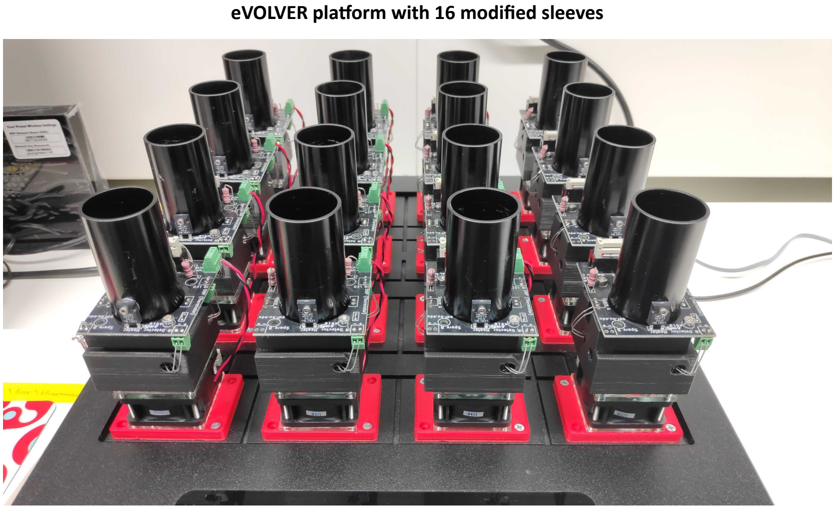


**Supplementary Fig. 3. Modified eVOLVER sleeve design.**
**(A)** Illustrated modifications resulted in stable and consistent OD measurement (as shown in Figure S6) along with cell culture light stimulation capability. **(B)** eVOLVER platform with 16 modified sleeves.

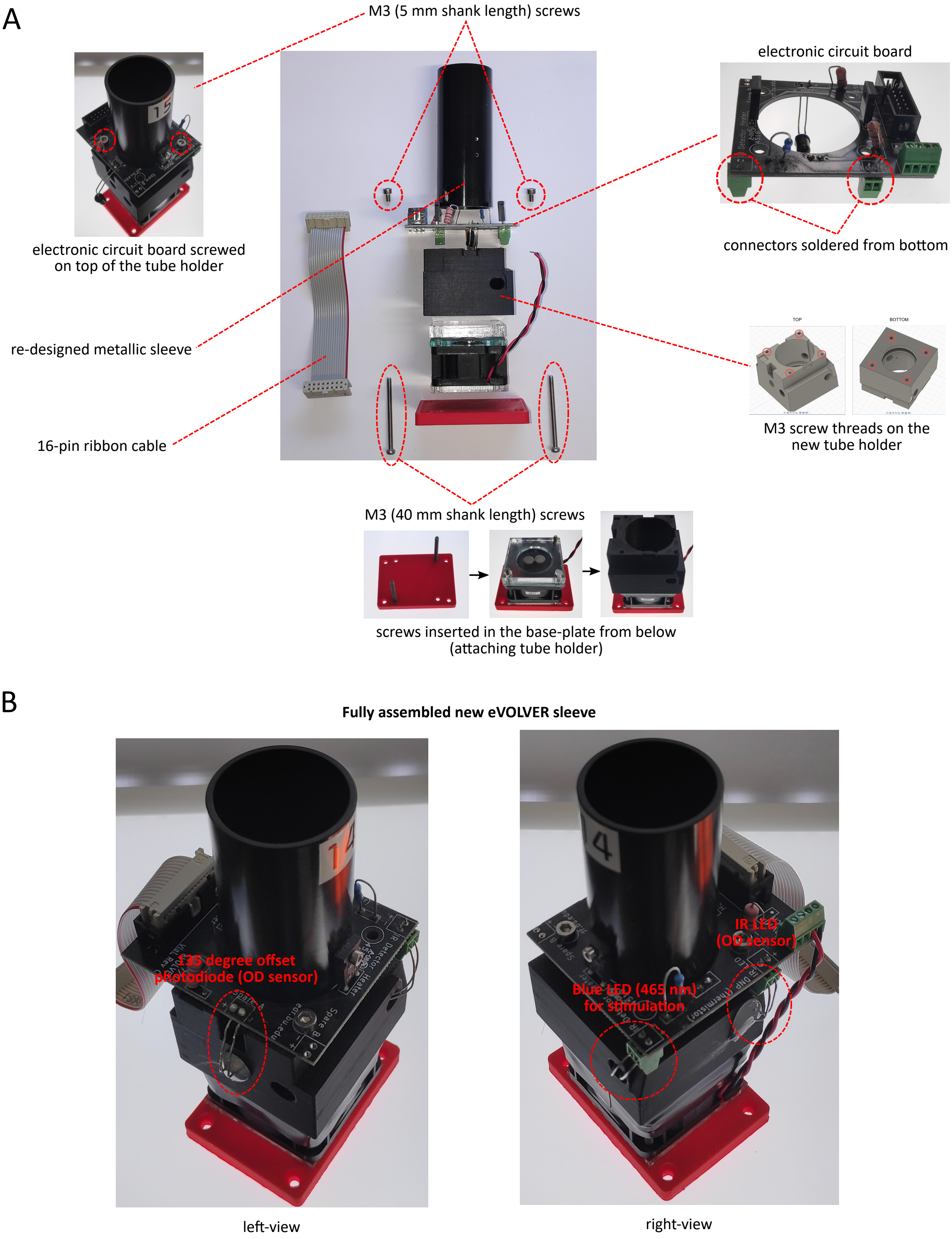


**Supplementary Fig. 4. Modified eVOLVER sleeve assembly.**
**(A)** Different components of the modified eVOLVER sleeve. **(B)** Fully assembled new eVOLVER sleeve with OD sensor (IR LED and photodiode) and blue LED for stimulation. This blue LED is connected to the electronic circuit board via a connector which was originally used for 90 degree offset photodiode in the default eVOLVER sleeve design.

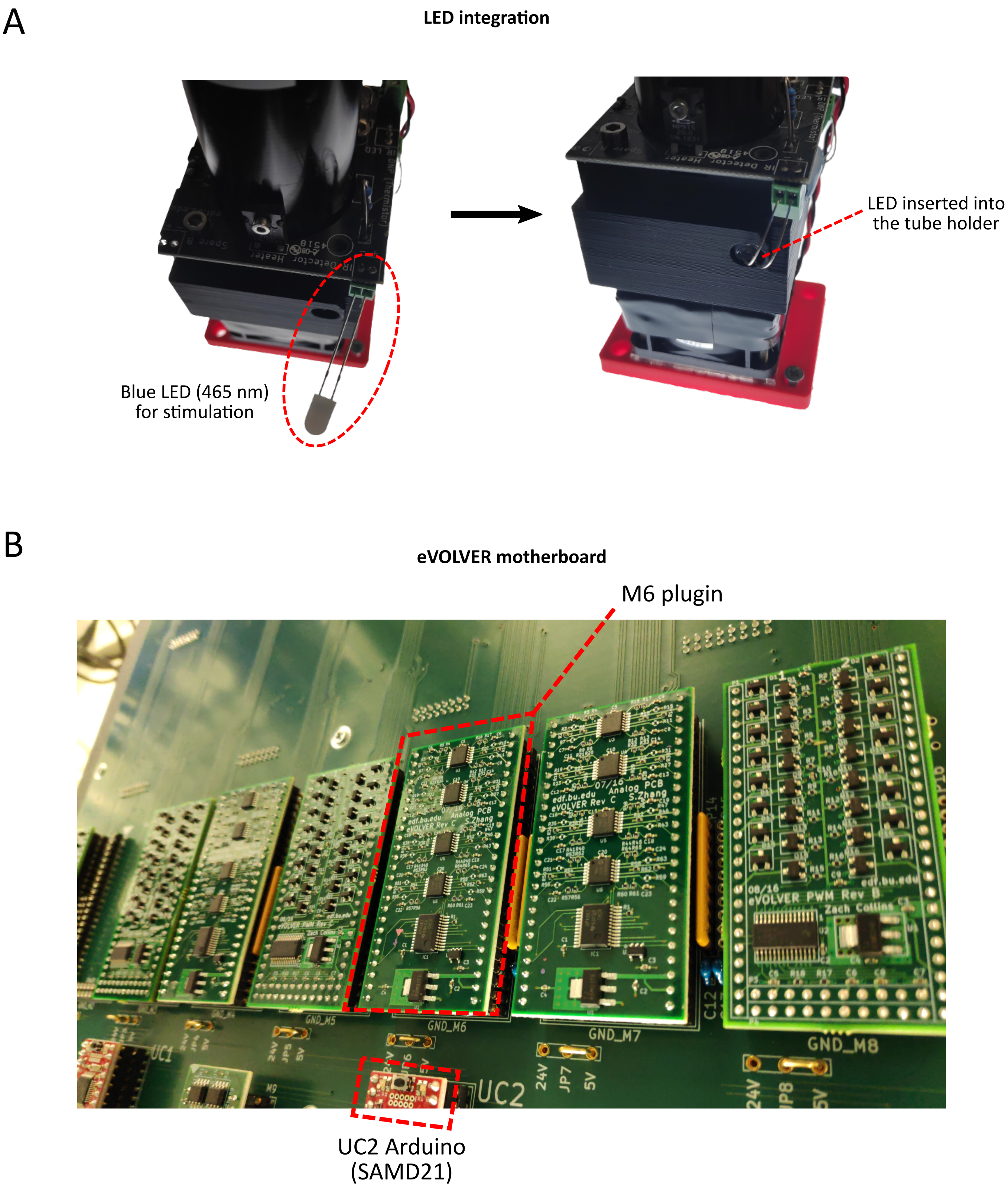


**Supplementary Fig. 5. LED integration on the modified eVOLVER sleeve.**
**(A)** LED for cell culture stimulation is connected to the sleeve electronic board at the 90 degree photodiode (IR detector) location. **(B)** To control and power this additional LED, M6 plugin on the eVOLVER motherboard is replaced with eVOLVER PWM board(1). UC2 SAMD21 arduino is re-programmed with suitable code accordingly, providing software control access for changing LED intensity during optogenetic experiments.

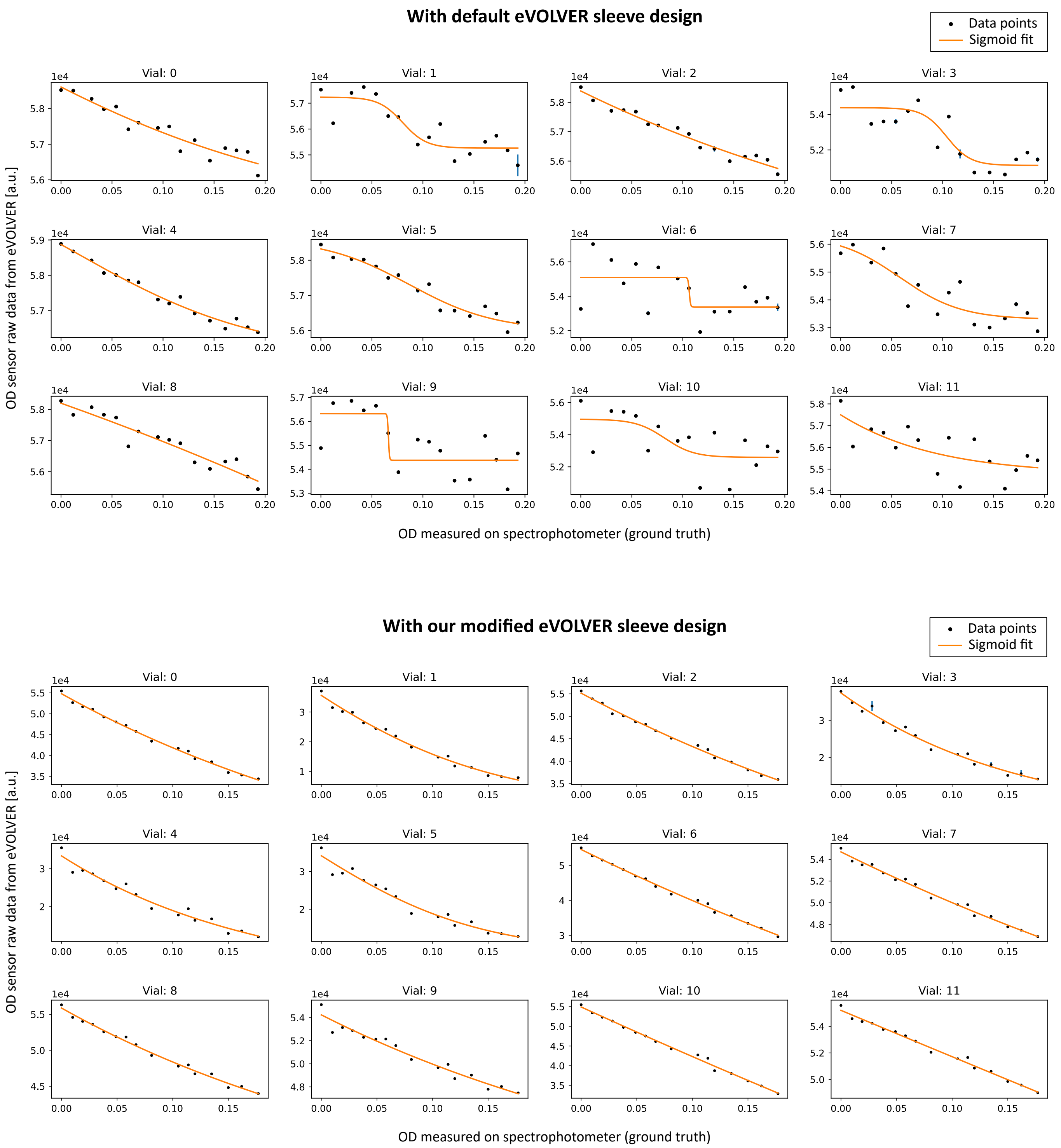


**Supplementary Fig. 6. Improved OD measurement with modified eVOLVER sleeve design**
As shown in Figure 3A **(Left)**, we re-designed the glass vial cap and the tube-holder with added O-rings to prevent wobbling of the culture vial inside the eVOLVER sleeve. This resulted in very stable and consistent OD sensor measurements compared to that with the default eVOLVER sleeve design.

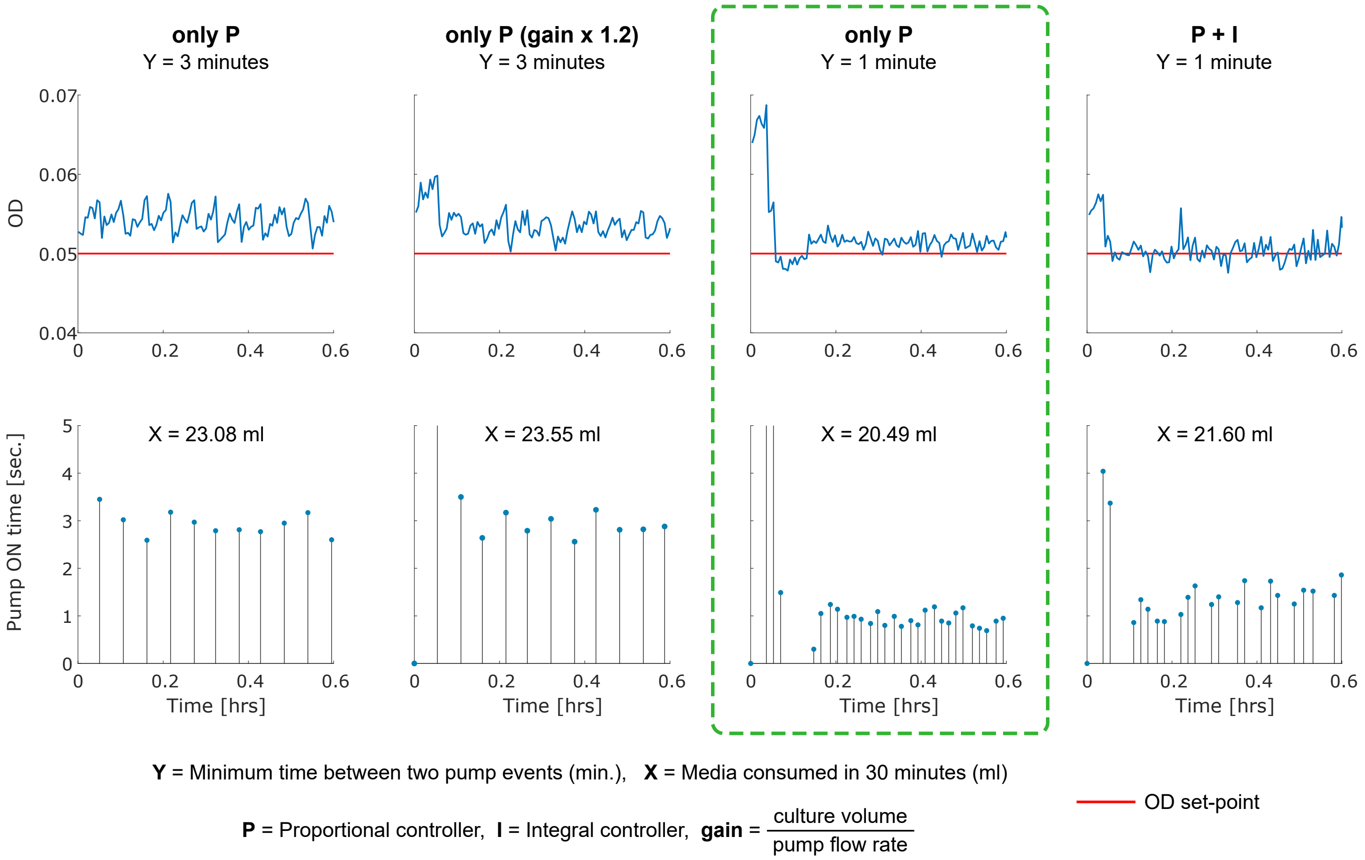


**Supplementary Fig. 7. OD regulation controller parameter tuning.**
We used the modified eVOLVER platform in turbidostat mode in order to maintain the cell culture density in a desired range in our experiments. This mode has an OD regulation feedback controller continuously running in the background. Based on OD measurements, this controller determines the duration for which the media pump should be switched ON in order to dilute the culture to a desired density. We performed multiple trial experiments with different OD regulation controller parameters. In all of our experiments in this study, we chose the controller parameters which exhibited better OD set-point tracking with less media consumption (highlighted with green dashed line box in this figure).

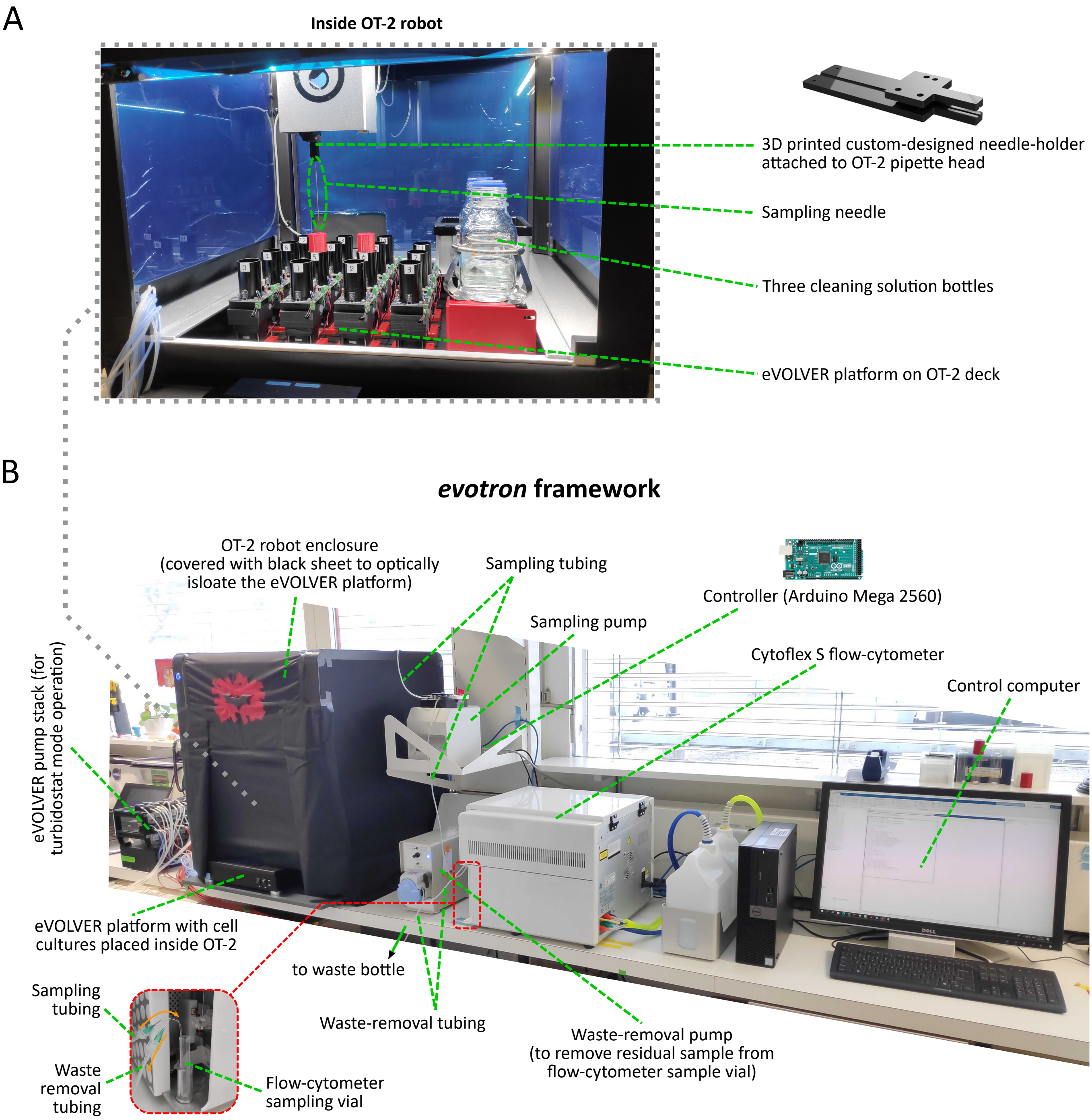


**Supplementary Fig. 8.** ***evotron*** **framework**
**(A)** Modfied eVOLVER platform placed inside Opentrons OT-2 robot **(B)** Complete *evotron* framework assembled on a lab bench.

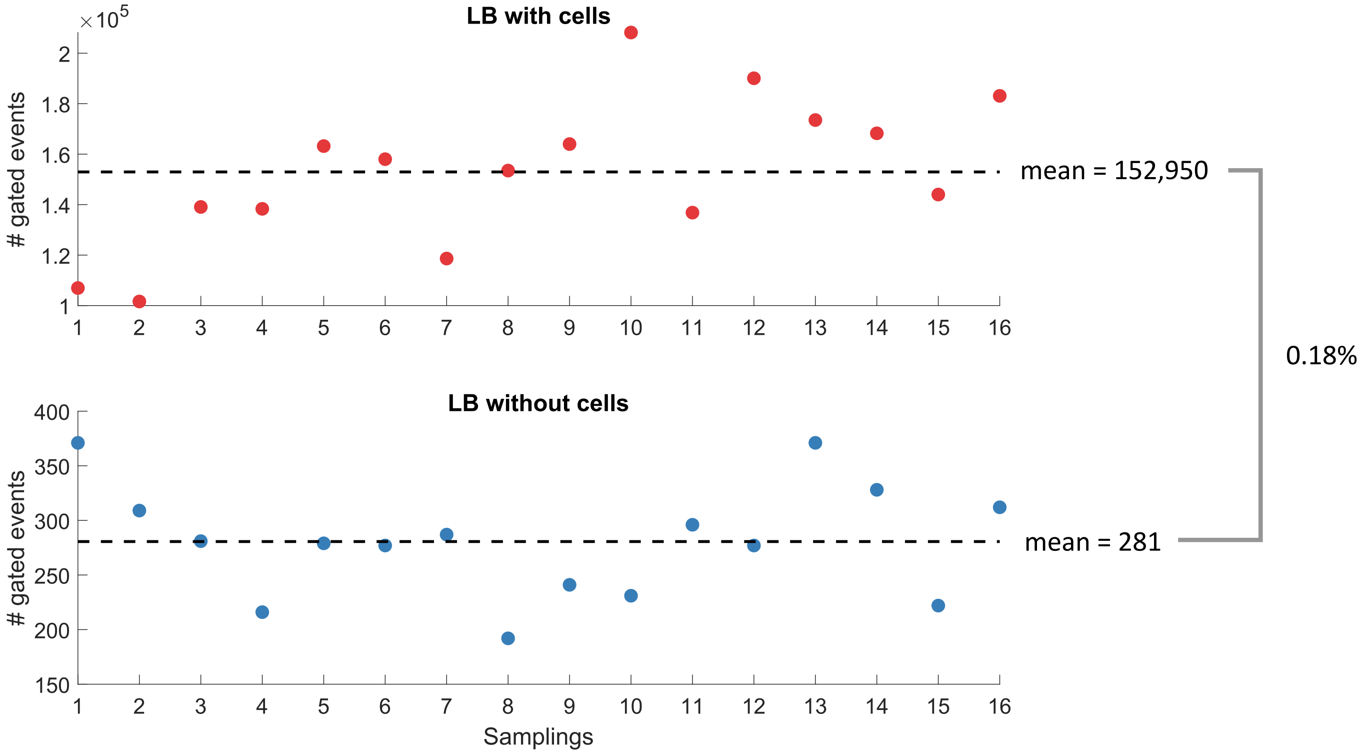


**Supplementary Fig. 9. Sampling with *evotron* platform exhibits negligible cross-contamination**
This figure shows the measured flow-cytometry gated events (indicating cell counts) in multiple automated samplings, with our *evotron* sampler, from two parallel cultures. One culture contained *E. coli* cells in LB media while the other contained sterile LB media without cells. Samples from these two cultures were taken in an alternate fashion with automated cleaning steps between successive sampling attempts. The number of gated events measured from the LB culture samples without cells displayed only background events with no increasing trend, even after multiple alternate sampling of LB culture with cells. Moreover, overnight incubation of the vial without cells after the 16th sampling cycle resulted in no bacterial growth, indicating that there had been no cross-contamination.

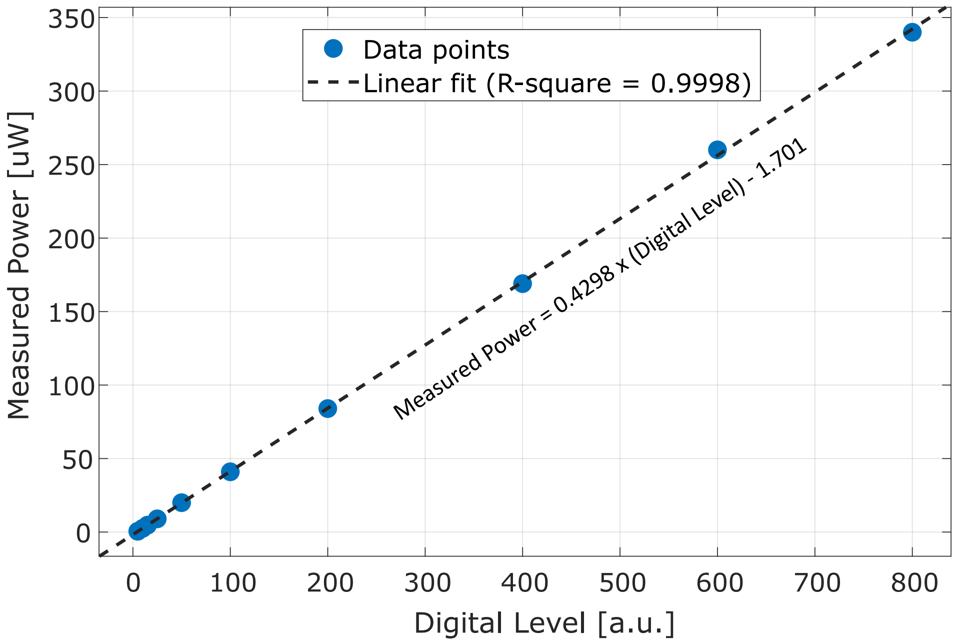


**Supplementary Fig. 10. LED Measured power vs Digital level**
We measured the blue LED (465nm) power inside the eVOLVER sleeve at different digital levels for characterization. Data shows a linear relation between the measured power and the applied digital level. LED power was measured with Nova power meter (PD300 sensor), Ophir Optronics Solutions Ltd.

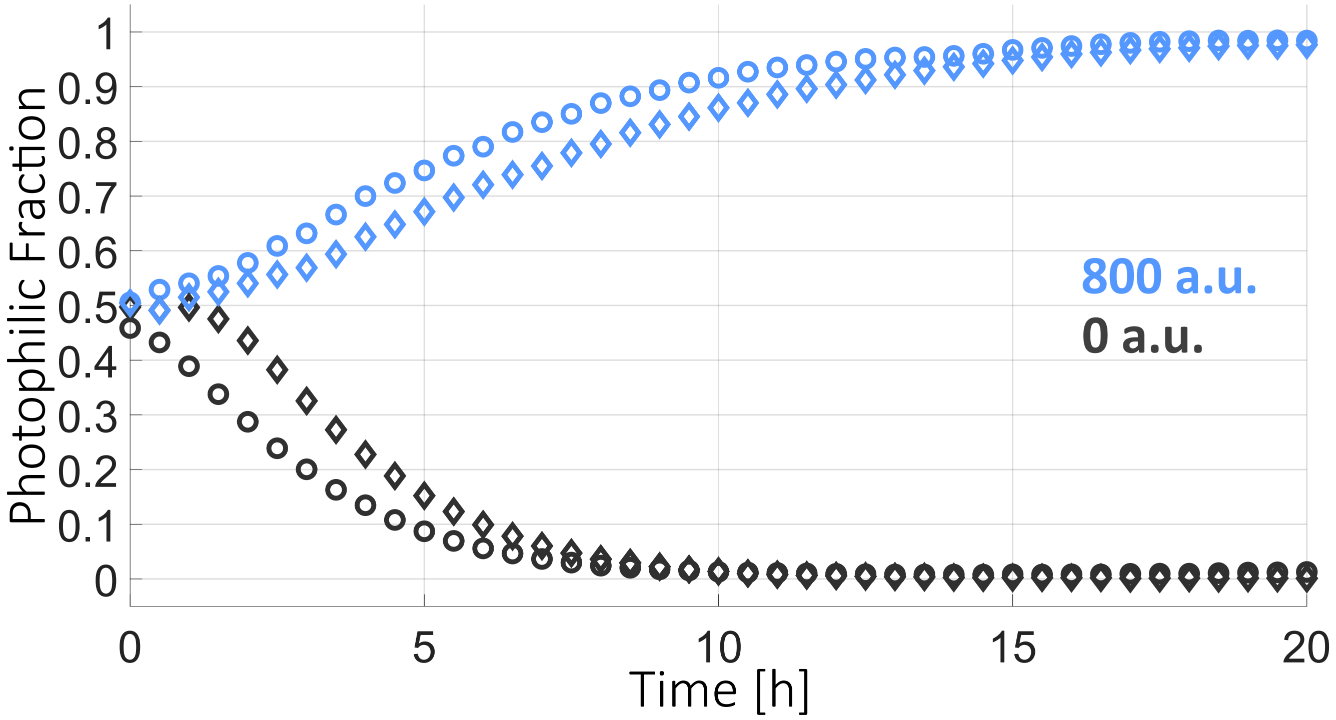


**Supplementary Fig. 11. Open-loop co-culture: replicates**
Biological replicates of the open-loop co-cultures with maximal light exposure or no light exposure shown in Figure 5B. Different symbols correspond to experiments carried out on different days.

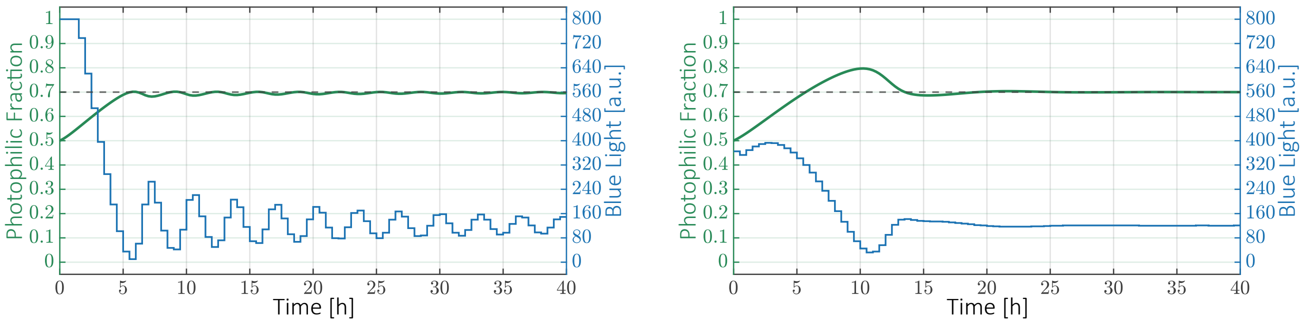


**Supplementary Fig. 12. Different sets of optimal PID gains**
The computational optimization procedure makes it possible to select gains that optimize different aspects of the dynamic behavior of the closed-loop system. **(Left)** The gains used in the closed-loop experiments produce a fast transient at the cost of low-amplitude oscillations at steady-state ($K_p = 5.9055 \cdot 10^3$, $K_i = 3.0382$, $K_d = 2.3427 \cdot 10^5$, $K_{bc} = 0.01 \cdot K_i$). **(Right)** An alternative set of gains results in no steady-state oscillations but after a longer transient phase with a large overshoot ($K_p = 1.5327 \cdot 10^3$, $K_i = 9.6743$, $K_d = 9.5689 \cdot 10^4$, $K_{bc} = 0.01 \cdot K_i$).

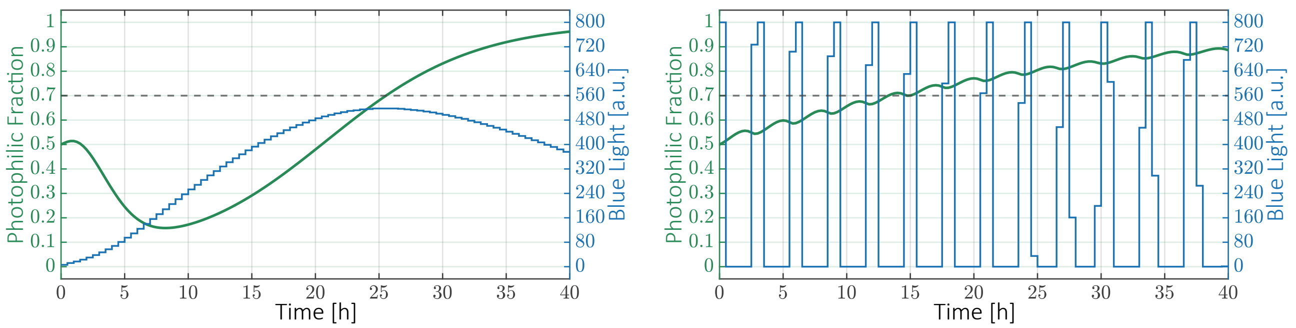


**Supplementary Fig. 13. Un-optimized PID gains lead to poor dynamic performance of the closed-loop system**
An arbitrary choice of gains lead to poor dynamic performance. Since the space of possible values for the gains is large, it is impractical to find a suitable set of gains through experimental trial-and-error. Model-guided optimization procedures can vastly reduce the number of experiments required to achieve optimal performance. **(Left)** $K_p = 1$, $K_i = 1$, $K_d = 1$, $K_{bc} = 1 \cdot K_i$. **(Right)** $K_p = 1 \cdot 10^5$, $K_i = 1$, $K_d = 1$, $K_{bc} = 1 \cdot K_i$.

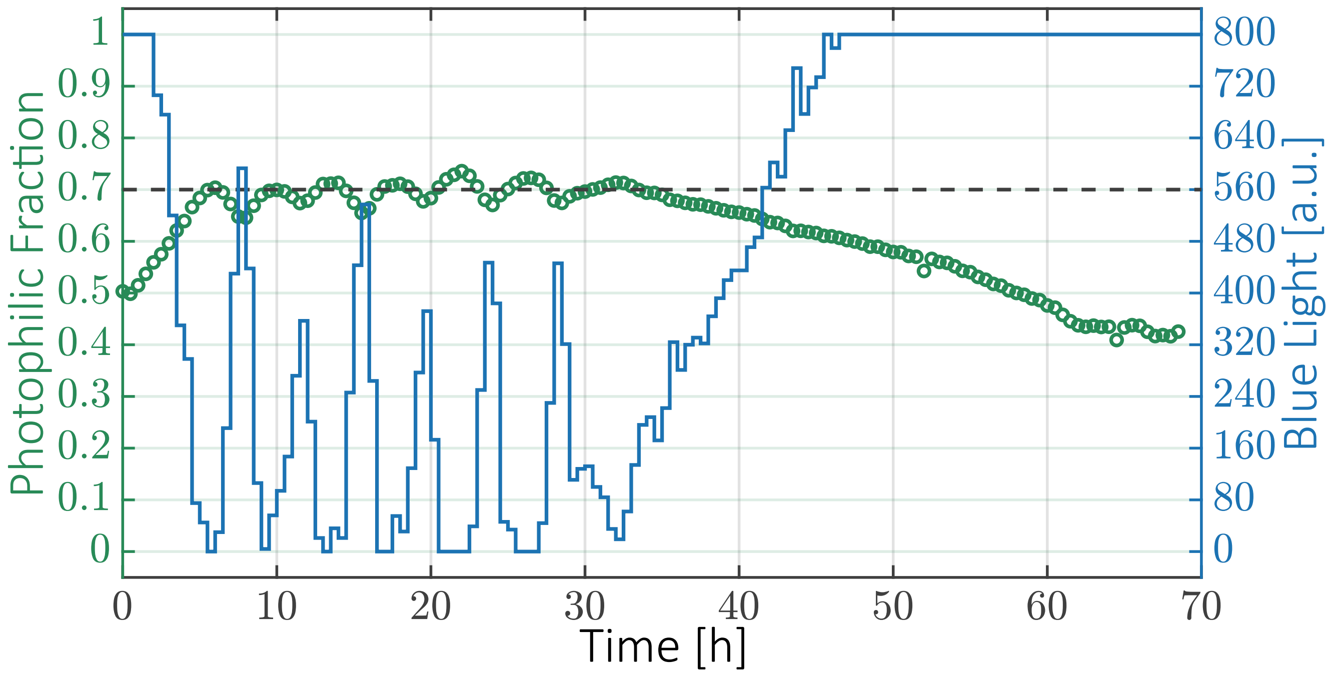


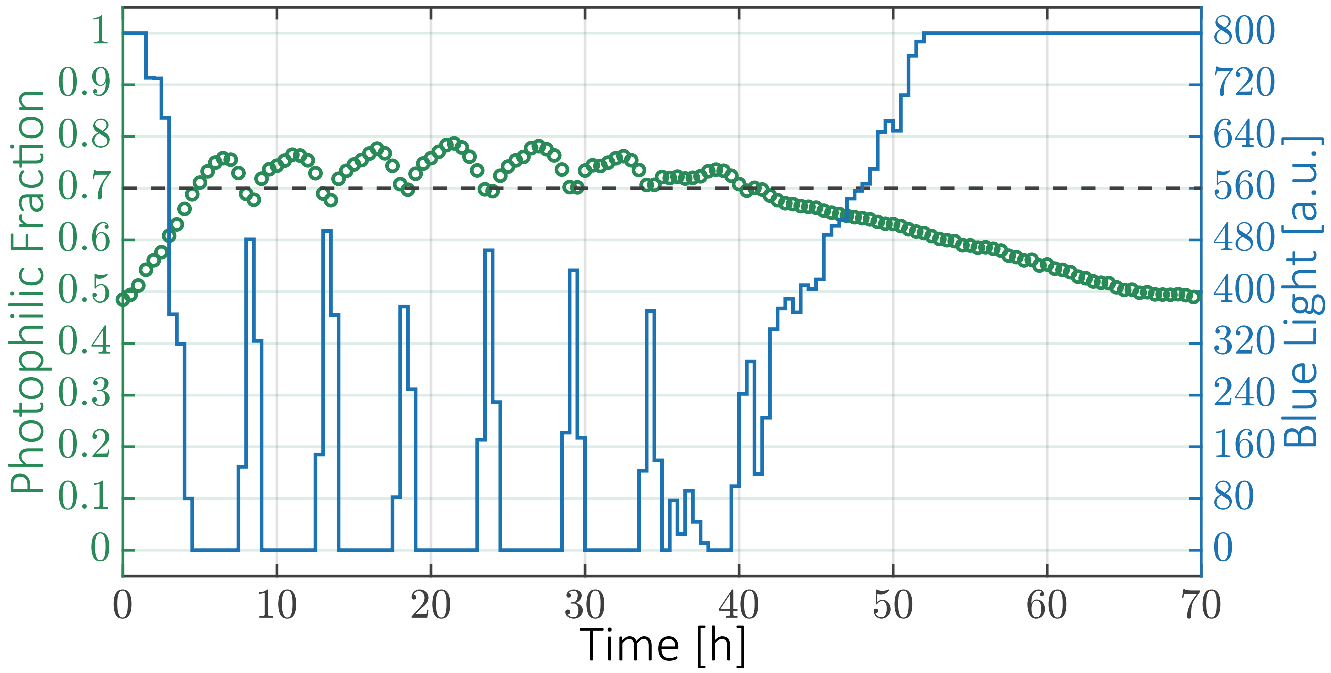


**Supplementary Fig. 14. High-setpoint: replicates**
Biological replicates of the high-setpoint closed-loop co-culture shown in Figure 6C.

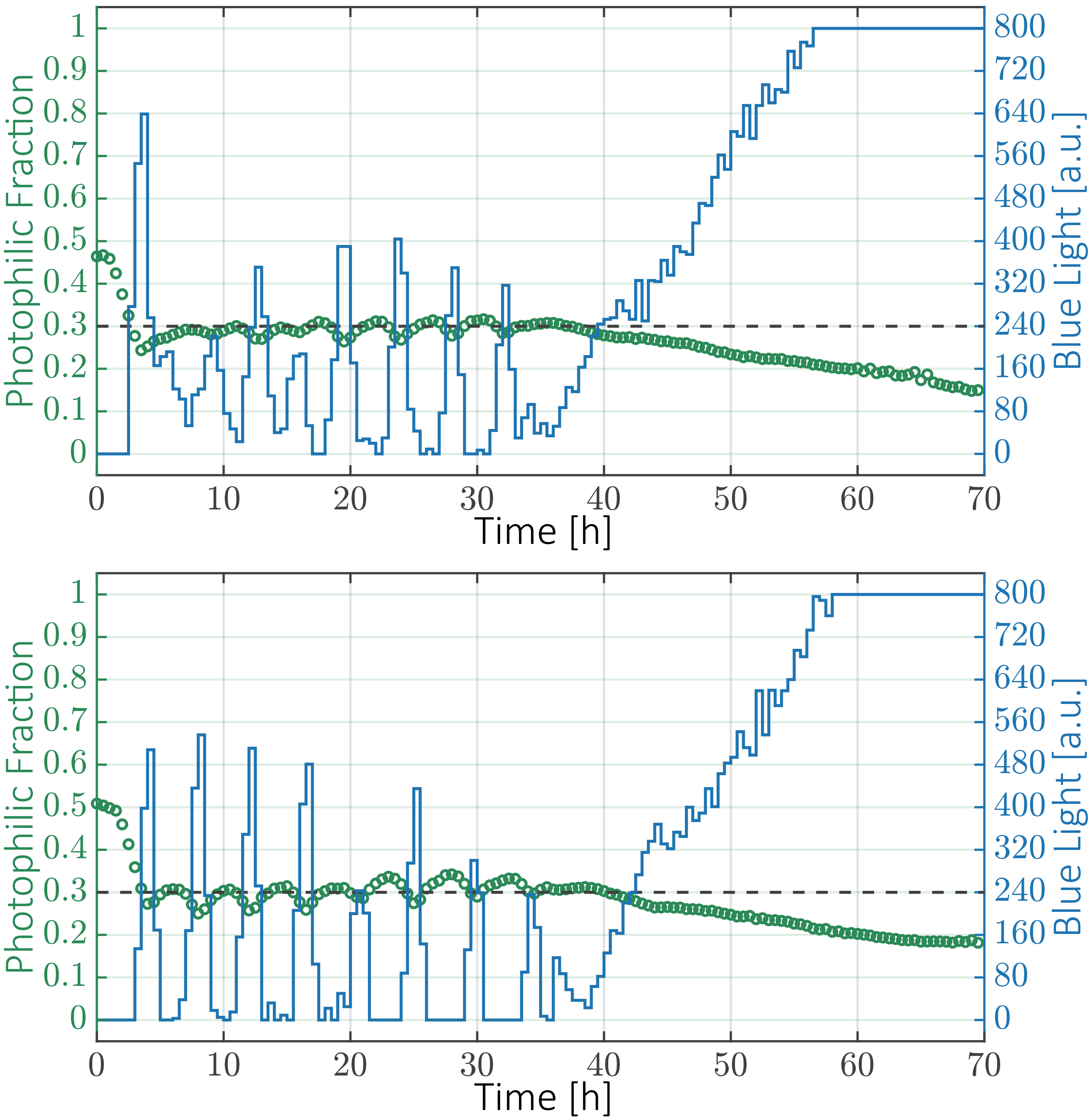


**Supplementary Fig. 15. Low-setpoint: replicates**
Biological replicates of the low-setpoint closed-loop co-culture shown in Figure 6D.

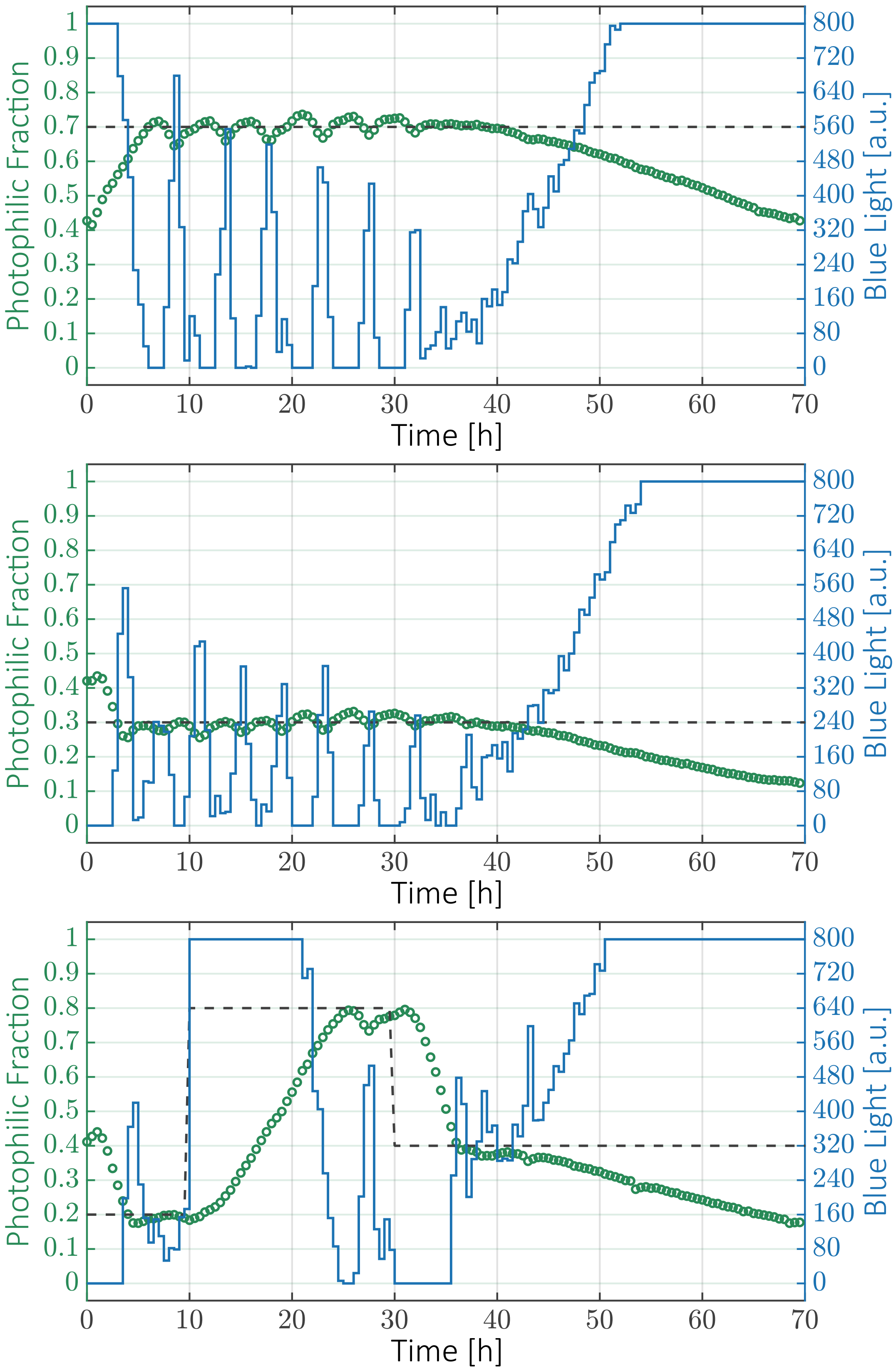


**Supplementary Fig. 16. Breakdown of stabilization after 40h**
After 40h we observe a continuous drift of the co-culture strain ratio in favor of the constitutive strain, probably reflecting the fixation of escape mutations in one of the strains or in both. The drift cannot be fully counteracted by the action of the controller, because change in growth rates due to mutations takes the closed-loop system out of the controllable regime. However, it can also be seen that the drift is much slower than what is observed in the absence of regulation (Figure 5), so that even 30h after the onset of the drift phase, the two strains still co-exist. We hypothesize that this is due to the fact that favorable mutations lead both strains to grow at the fastest attainable rate, which will be similar in both cases due to the low metabolic burden imposed by the growth control circuit in the photophilic strain.

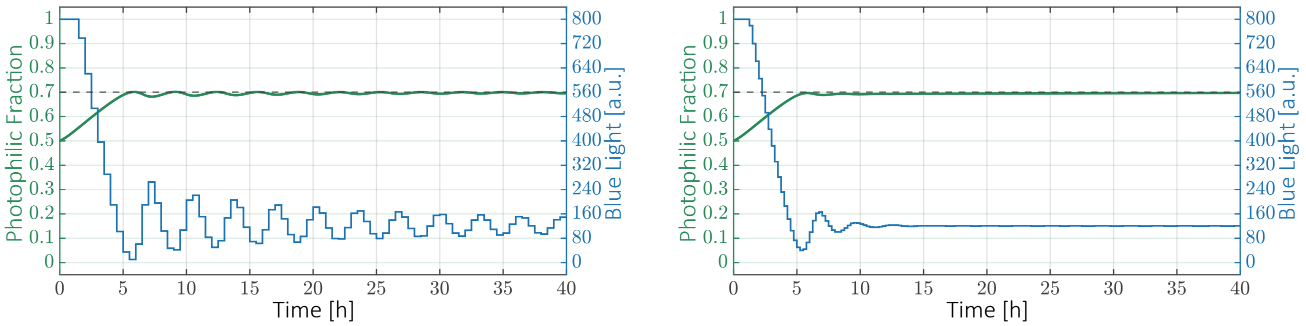


**Supplementary Fig. 17. Sampling frequency and stability**
The mathematical model suggests that the low-amplitude oscillations at steady-state observed in our closed-loop experiments (6) could be avoided if the sampling frequency is increased. **(Left)** Sampling every 30 minutes. **(Right)** Sampling every 15 minutes.

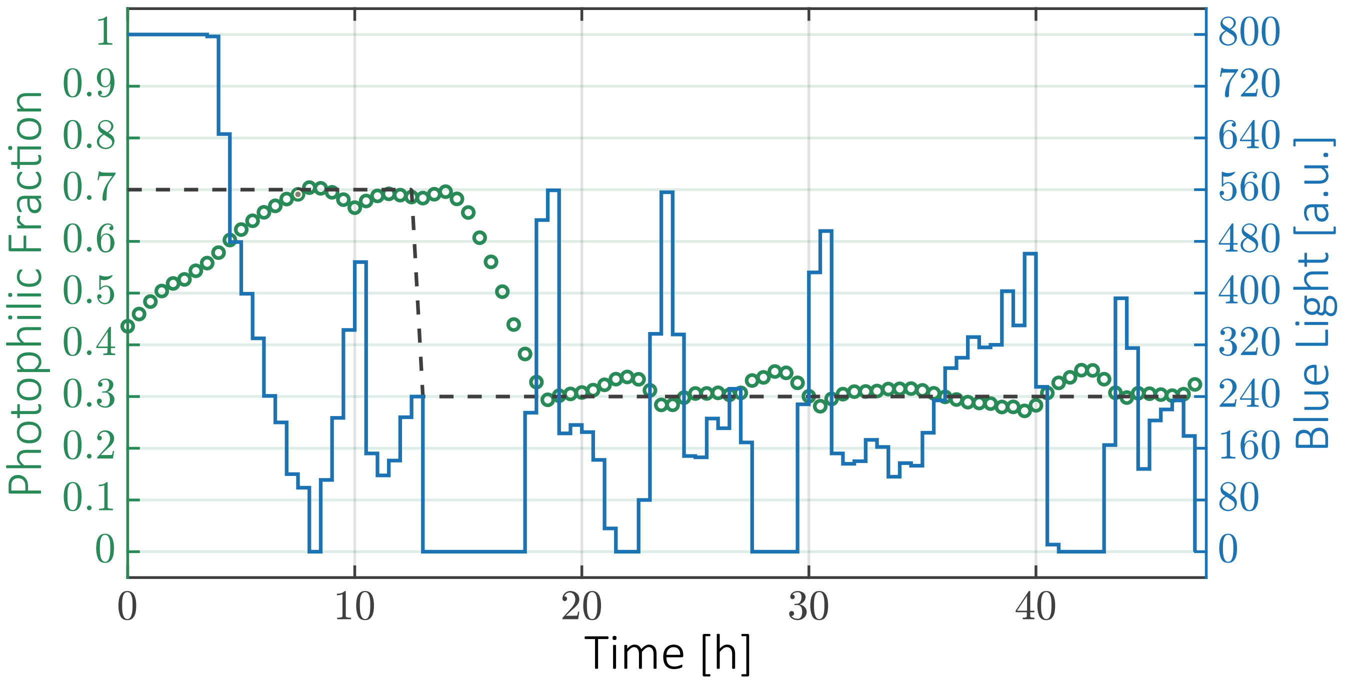


**Supplementary Fig. 18. Setpoint tracking**
As shown in Figure 6, the closed-loop co-culture can be forced to track a dynamically changing setpoint. In this case, the objective was to set the strain ratio to 0.7 during the first 13h and then to change it to 0.3 for the remainder of the experiment.

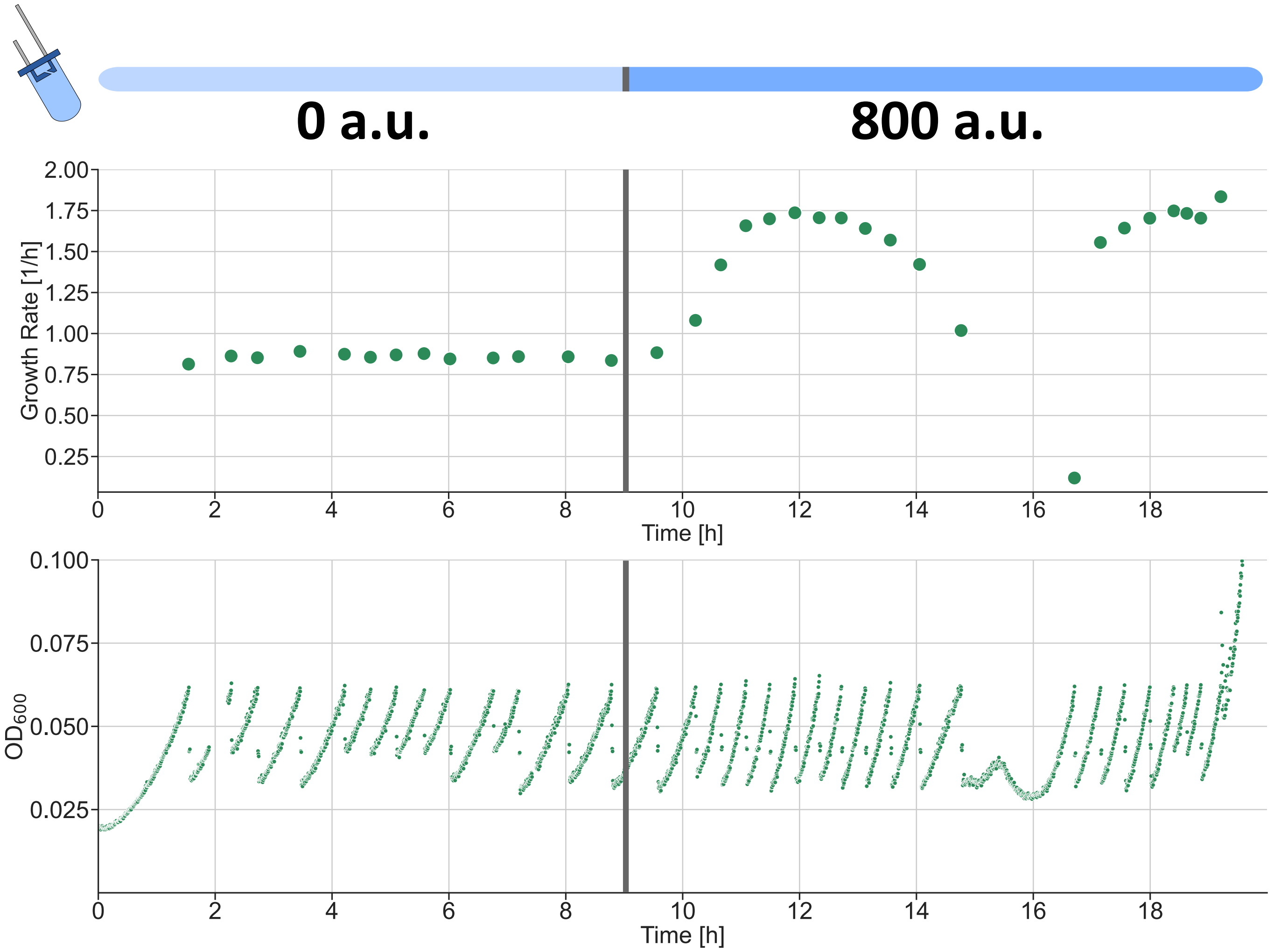


**Supplementary Fig. 19. Growth arrest on *evotron* platform**

After the *evotron* platform was assembled, we began observing a spontaneous, reversible growth arrest in the turbidostat cultures happening after 10-15h. The growth arrest happened at earlier times when the cells were grown with maximal light from the beginning, suggesting that the metabolic state of the cells influenced the timing of the arrest. Since this platform consists of a closed opentrons OT-2 robot (tightly covered with black foil to avoid ambient light inside) with a modified eVOLVER platorm replacing its deck (Figure S8), we hypothesize that this growth arrest was caused by insufficient aeration within the covered OT-2 enclosure. Addition of an external source of pressurized air into the chamber restored normal growth (Figure S20) and the growth defect was never observed when the air supply was running.

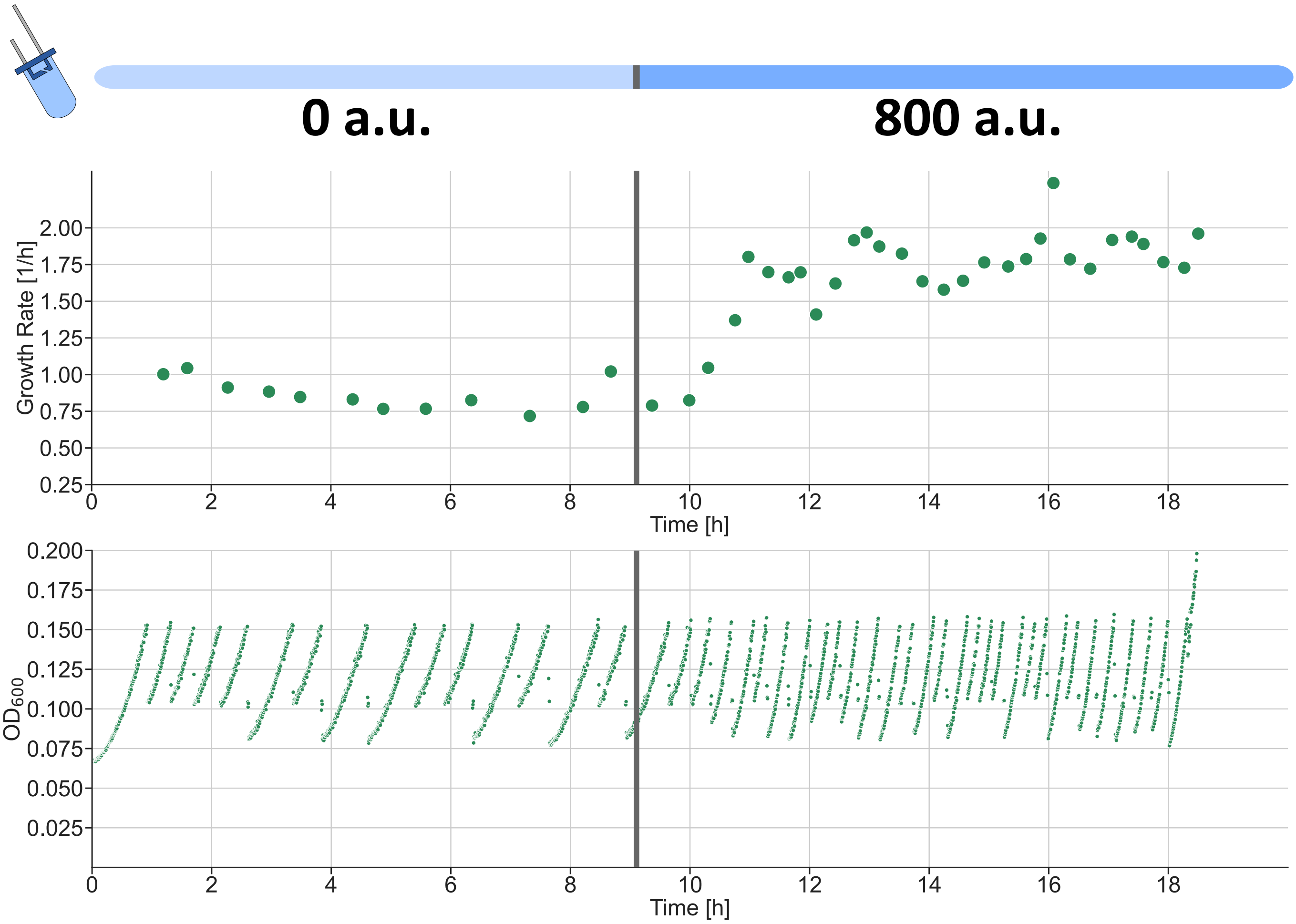


**Supplementary Fig. 20. Aeration restores normal growth on *evotron* platform**
Addition of an external source of pressurized air (4 bar) into the covered OT-2 enclosure restored normal growth in the turbidostat cultures. The data in this figure corresponds to an experiment analogous to the one shown in Figure S19, but where the external supply of air was active throughout the experiment. The OD tolerance range of the turbidostat is different between the two experiments, but we observed that this was not the cause of the growth arrest.

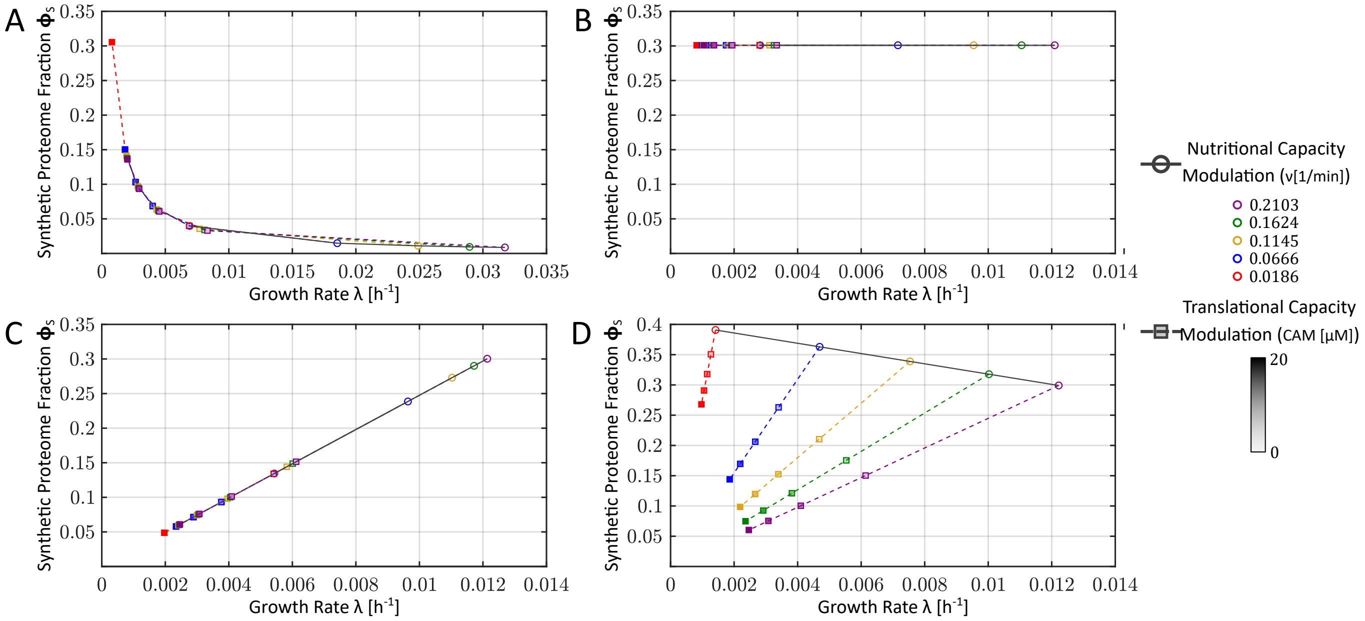


**Supplementary Fig. 21. Model predictions for the dependency of the expression of a constitutive gene on growth rate** The proteome fraction occupied by the constitutive protein is plotted against the growth rate. In the model, growth is varied by two independent mechanisms, changing the quality of nutrients (nutritional capacity, different colors) or by the presence of sublethal concentrations of chloramphenicol (translational capacity, squares, increasing opacity denotes increasing CAM concentrations). **(A)** Conventional non-host-aware model. **(B)** Model that includes translational resources. **(C)** Model that includes both transcriptional and translational resources. **(D)** Model that includes both transcriptional and translational resources, as well as an explicit dependency on the nutritional capacity $\nu$.

| Parameter | Description | Value | Unit | Source |
|---|---|---|---|---|
| $\hat{\alpha}_T$ | Effective T7-monomer production rate | $7.2 \cdot 10^{-3}$ | nM | $\star$ |
| $\hat{\alpha}_C$ | Effective CAT production rate | $6.467 \cdot 10^{-6}$ | $\mathrm{nM\,min^{-1}}$ | $\star$ |
| $K_G$ | $K_m$ of T7-promoter activation | 57.1585 | nM | $\star$ |
| $h_{ON}^{min}$ | Minimal dimerization rate | $2.0202 \cdot 10^{-7}$ | $\mathrm{nM^{-1}\,min^{-1}}$ | $\star$ |
| $h_{ON}^{max}$ | Maximal dimerization rate | $2.0020 \cdot 10^{-5}$ | $\mathrm{nM^{-1}\,min^{-1}}$ | $\star$ |
| $K_L$ | Light-dependent dimerization's $K_m$ | $1.9851 \cdot 10^{3}$ | a.u.$*$ | $\star$ |
| $n_L$ | Light-dependent dimerization's Hill coefficient | 1.3548 | | $\star$ |
| $n_G$ | Hill coefficient of T7-promoter activation | 1.5557 | | $\star$ |
| $h_C$ | Hill-exponent of CAM degradation | 1.5388 | | $\star$ |
| $\gamma_{\text{mol2fluo}}$ | Conversion factor between molecule numbers and flow-cytometry arbitrary mCherry fluorescence units | 0.2549 | $\mathrm{nM^{-1}}$ a.u. $**$ | $\star$ |
| $L_0$ | Effective light-intensity corresponding to ambient light during pre-culturing | 196.3930 | a.u.$*$ | $\star$ |
| $N_p$ | p15A plasmid copy number (CAT gene) | 10 | | (14) BNID: 105307 |
| $K_D$ | $K_m$ of CAM-ribosome binding reaction | 1300 | nM | (7) |
| $\hat{K}_C$ | $K_m$ of CAM-ribosome binding reaction | 0.3333 | nM min | (7) |
| $\kappa$ | CAM diffusion rate | 90 | $\mathrm{min^{-1}}$ | (7) |
| $n_r$ | effective ribosomal unit length | 12221 | aa | Estimated from (2) |
| $n_T$ | T7-monomer length | 597 | aa | ‡ |
| $n_C$ | CAT length | 219 | aa | |
| $g_0$ | Average translation rate per ribosome | 0.0987 | $\mathrm{min^{-1}}$ | Estimated from (2) |
| $\Phi_R^{max}$ | Maximal ribosomal proteome fraction | 0.5470 | | (2) |
| $\Phi_{R_0}$ | Proteome fraction of inactive ribosomes | 0.0660 | | (2) |
| $\rho_{cell}$ | Cell density (conversion factor between proteome fractions and molecules per cell | $2 \cdot 10^{9}$ | $\mathrm{aa\,fL^{-1}}$ | (5) |
| $\nu$ | Nutrient capacity of medium (LB) | 0.1921 | $\mathrm{min^{-1}}$ | † |
| $A_E$ | External concentration of CAM | $10.5 \cdot 10^{3}$ | nM | |

**Supplementary Table 1. Parameter values used for simulations of photophilic strain and co-culture dynamics** $\star$Obtained from fit to the photophilic strain's dynamic response. †Obtained from manual fitting to the growth rate of the photophilic strain in the absence of CAM. ‡Average between the length of the two T7 split units(15). $*$Arbitrary units of light intensity. $**$Arbitrary units of flow-cytometry mCherry fluorescence.
